# PAF1 facilitates RNA polymerase II ubiquitination by the Elongin A complex through phosphorylation by CDK12

**DOI:** 10.1101/2020.09.17.297960

**Authors:** Gabriel Sanchez, Jérôme Barbier, Céline Elie, Rosemary Kiernan, Sylvie Rouquier

## Abstract

The conserved Polymerase-Associated Factor 1 complex (PAF1C) regulates all stages of the RNA polymerase II (RNAPII) transcription cycle from the promoter to the 3’ end formation site of mRNA encoding genes and has been linked to numerous transcription related processes. Here, we show that PAF1 interacts with Elongin A, a transcription elongation factor as well as a component of a cullin-RING ligase that targets stalled RNAPII for ubiquitination and proteasome-dependent degradation in response to DNA damage or other stresses. We show that, in absence of any induced stress, PAF1 physically interacts with the E3 ubiquitin ligase form of the Elongin A complex and facilitates ubiquitination of RNAPII. We demonstrate that this ubiquitination is dependent of the Ser2 phosphorylation of the RNAPII carboxy-terminal domain (CTD) by CDK12. Our findings highlight a novel unexpected role of PAF1-CDK12 in RNAPII transcription cycle, raising the possibility that the Elongin A ubiquitin ligase plays a role in normal transcription process, and suggest a transcription surveillance mechanism ready to degrade RNAPII if needed.

## INTRODUCTION

Transcription elongation is a very dynamic and discontinuous process involving frequent and prolonged arrests of RNA polymerase II (RNAPII) following different events including DNA damage, topological constraints and nucleotide depletion, among others. Many factors contribute to the processivity of RNAPII helping to manage these transitory pauses and stalls (Wilson et al., 2013). However, if the RNAPII is unable to resume elongation, this transcriptional pause can evolve towards a transcriptional stop and the polymerase becomes irreversibly arrested. The stalled RNAPII molecule must be removed from the DNA template to allow repair and subsequent transcription. Transcription-stalled RNAPII results in the polyubiquitination and degradation by the proteasome of the largest subunit of RNAPII, RPB1 (Bregman et al., 1996; Gregersen and Svejstrup, 2018; Mitsui and Sharp, 1999; Woudstra et al., 2002). Although originally identified as a response to DNA damage, RPB1 polyubiquitination and degradation occurs under a number of conditions leading to transcriptional arrest (Somesh et al., 2005; Wilson et al., 2013). Removal of arrested RNAPII complexes is accomplished, in part, through ubiquitination of RNAPII *via* a Cullin-RING E3 ligase (CRL) with Elongin A (EloA or TCEB3) as its substrate recognition subunit. The Elongin complex contains EloA, and two smaller regulatory subunits Elongin B (EloB or TCEB2) and Elongin C (EloC or TCEB1). It interacts with the phosphorylated form of RNAPII C-terminal domain (CTD) and stimulates transcription elongation ((Conaway and Conaway, 2019) and references therein). However, along with Cullin family member CUL5 and RING finger protein RBX2, this complex drives degradation of stalled RNAPII by the proteasome (Harreman et al., 2009; Kawauchi et al., 2013; Yasukawa et al., 2008). The assembly of the ligase is tightly regulated and rapidly induced not only with a large variety of DNA-damaging agents but also with drugs that induce RNAPII stalling (Weems et al., 2015). RNAPII ubiquitination is a two-step process that requires first a monoubiquitination of RPB1 by NEDD4/Rsp5, followed by the polyubiquitination by the Elongin A ubiquitin ligase (Harreman et al., 2009; Yasukawa et al., 2008).

Since the degraded RNAPII is hyper-phosphorylated, it suggests that the elongating form of RPB1 is targeted by ubiquitin ligase complexes (Mitsui and Sharp, 1999). CTD Serine 2 (Ser2) phosphorylation has been associated with productive transcriptional elongation (pause-release of the RNAPII) and many aspects of mRNA processing, such as splicing and 3’ end processing (Buratowski, 2005; Heidemann et al., 2013). Although cyclin-dependent kinase 9 (CDK9) has long been thought to be the predominant Ser2 kinase, studies have identified CDK12 as a major CTD Ser2 kinase (Bartkowiak et al., 2010; Cheng et al., 2012; Czudnochowski et al., 2012). In addition, CDK12 is involved in transcription termination. Polyadenylation-coupled Ser2 phosphorylation by CDK12 is required for recruitment of cleavage and polyadenylation factors (Davidson et al., 2014).

The PAF1 complex (PAF1C) in humans is comprised of six protein subunits: PAF1, CDC73, CTR9, LEO1, RTF1 and WDR61 (Zhu et al., 2005), and was originally identified in yeast as a RNAPII-interacting protein complex (Mueller and Jaehning, 2002). PAF1C presents no enzymatic activity and has been implicated in the regulation of different steps of transcription: promoter-proximal pausing (Chen et al., 2015; Yu et al., 2015), transcription elongation, recruitment and activation of histone modifiers, mRNA 3’-end formation (for review (Van Oss et al., 2017) and references therein). PAF1C is required for H2B ubiquitylation and downstream H3K4 and H3K79 methylation both in yeast (Krogan et al., 2003; Ng et al., 2003; Wood et al., 2003) and humans (Kim et al., 2009; Zhu et al., 2005). It has been shown that PAF1C triggers CDK12 recruitment to allow the release of paused RNAPII (Yu et al., 2015). PAF1C is also involved in the resolution of transcription-replication conflicts (Poli et al., 2016) and was shown to modulate elongation rate of the RNAPII (Hou et al., 2019).

To further explore the PAF1C function in transcription regulation, we analyzed its interactome by tandem affinity purification followed by mass spectrometry. We identified the Elongin complex and confirmed it is a partner of the PAF1C and we also showed that PAF1 interacts with the Elongin A E3 ubiquitin ligase. We demonstrate that PAF1 through the recruitment of CDK12 and the subsequent phosphorylation of the CTD facilitates RNAPII ubiquitination under basal conditions. Our study reveals an unexpected role of PAF1-CDK12 in regulating RNAPII degradation during transcription elongation and suggests a surveillance mechanism that senses stalled RNAPII, ready to facilitate RNAPII degradation when needed.

## RESULTS

### PAF1 nuclear interactome analysis

To get more insights into PAF1C functions during transcription, we purified PAF1 and its associated partners. We cloned full-length PAF1 fused to FLAG and HA epitope tags at its N terminus. HeLa S3 cells expressing this construct were used to prepare Dignam nuclear extracts that were subjected to tandem immunoaffinity chromatography with anti-FLAG and anti-HA antibodies (Figure S1A). PAF1-associated partners were visualized by silver-staining following SDS-PAGE (Figure 1A) and identified by tandem mass spectrometry (Figure 1B and Table S1). Among the 182 proteins identified, as expected subunits of the PAF1C, including CTR9, LEO1, CDC73, WDR61 (SKI8), were recovered as well as known PAF1C interacting partners such as SUPT16H, SSRP1 or WDR82 (Figures 1A, 1B and Table S1). Analyses of biological processes (Gene ontology analysis, Figure 1C and Table S2) and pathways (KEGG and Reactome analysis, Figure S1B and Table S3) showed that the majority of PAF1 interactants are implicated in gene regulation and DNA conformation changes. Interestingly, the two subunits of the Elongin A complex, EloA and EloB were recovered among PAF1 interactants (Figure 1B and Table S1).

**Figure 1.**
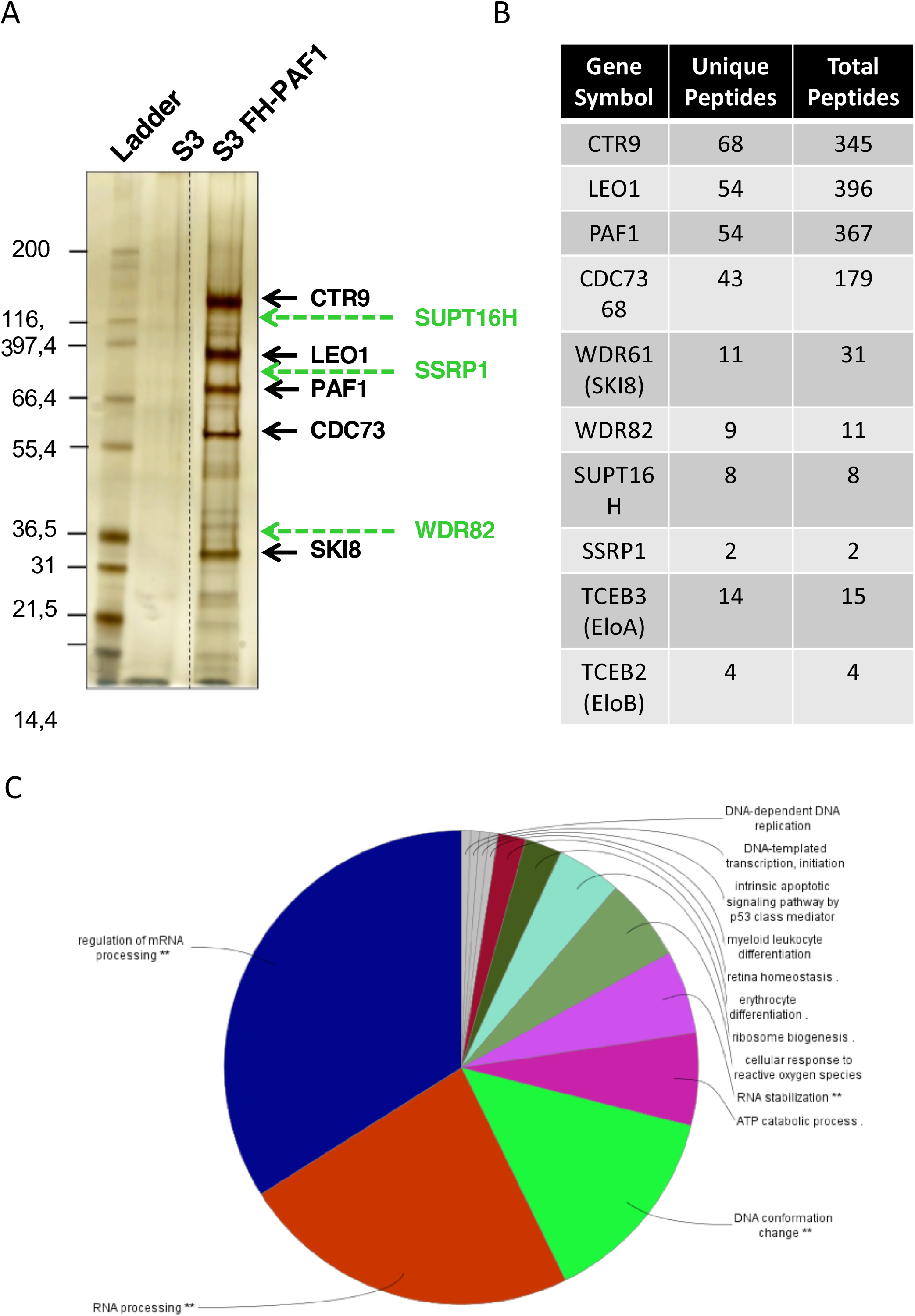
Identification of PAF1 partners. **(A)** SDS-PAGE analysis followed by silver staining of eluates of tandem affinity purified Dignam nuclear extracts from HeLa S3 mock cells or stably expressing FLAG-HA-PAF1 (FH-PAF1). Subunits of the PAF1C are shown in black and major known partners of the PAF1C are indicated in green. **(B)** Number of unique peptides and total peptides of FH-PAF1 partners identified by tandem mass spectrometry (MS/MS). The complete list of partners is listed in Table S1. **(C)** Pie chart of biological processes (GO) of proteins identified by mass spectrometry. The proteins involved are listed in Table S2.

### The Elongin ligase complex interacts with PAF1

Endogenous interactions between PAF1 and Elongin A complex were supported by co-immunoprecipitation (co-IP) analysis in HCT116 cells by probing EloA or EloB on immunoprecipitations performed using antibodies against either PAF1 or LEO1, two subunits of PAF1C (Figure 2A). Reverse immunoprecipitations of endogenous proteins using EloA or EloB antibodies as indicated (Figure 2B) or overexpressed EloA protein (Figure S2A) also probed positively with PAF1 antibody. Since PAF1 and Elongin A complexes are both involved in transcription elongation, this association was not so surprising. However, the Elongin heterotrimer assembles with CUL5 and RING finger protein Rbx2 to form an E3 ubiquitin ligase that targets the RPB1 subunit of RNAPII for ubiquitination and degradation by the proteasome in cells subjected to different stresses (Weems et al., 2015). To get further insight on the functional significance of the PAF1-Elongin complex interaction, we decided to test whether the E3 ligase activity might also be involved. We performed co-IPs using either PAF1 or LEO1 and detected CUL5, suggesting that PAF1 complex interacts with the Elongin A E3 ligase. These co-IPs were performed using CUL5-transfected cells as the endogenous CUL5 could not be retrieved even in an EloA immunoprecipitation using various antibodies (data not shown) (Figure 2C).

**Figure 2.**
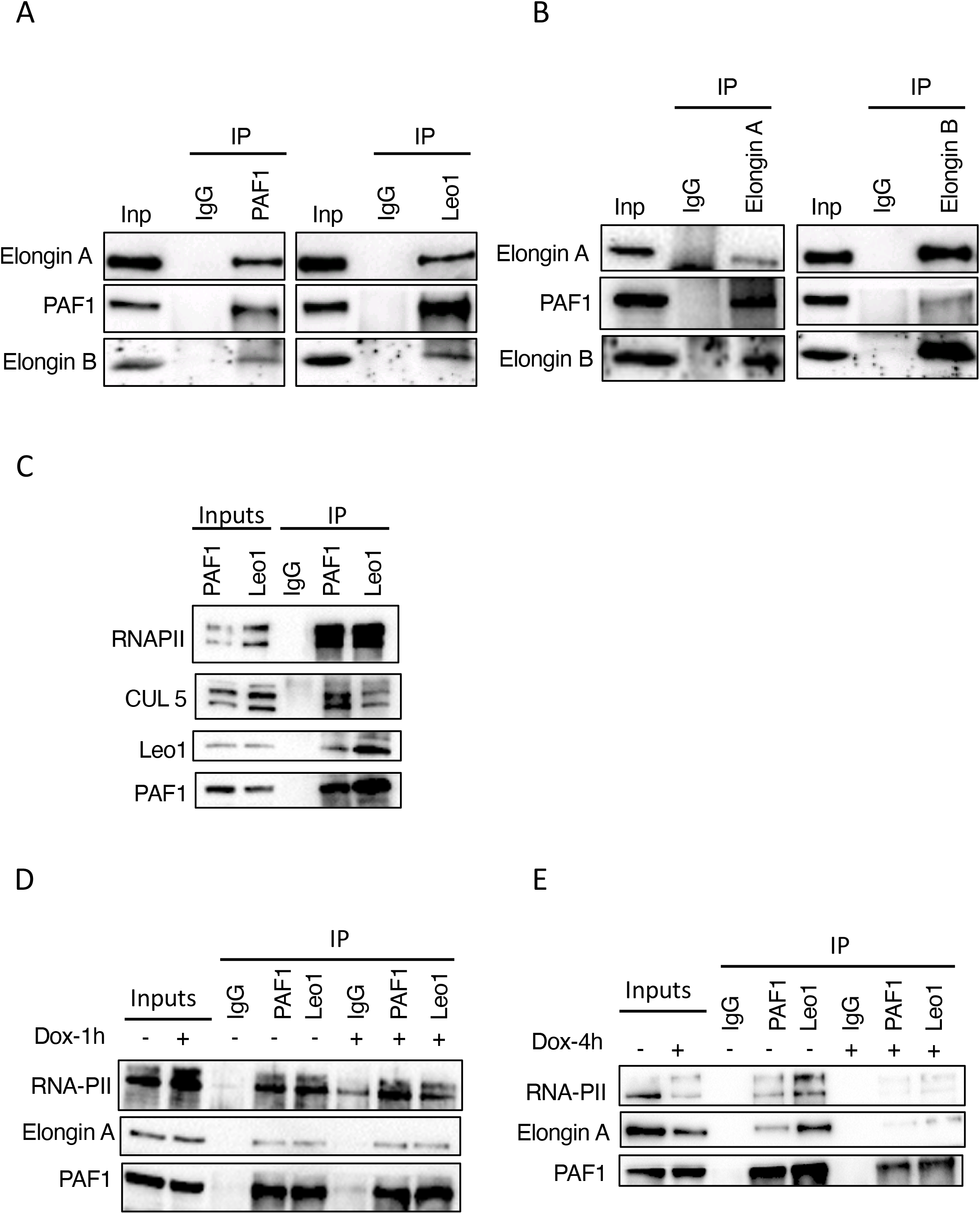
PAF1 interacts with the Elongin A ligase complex. **(A-E)** Proteins from HCT116 nuclear extracts were immunoprecipitated with the indicated antibodies. Nuclear extracts (Inputs 5%) and immunoprecipitates were separated by SDS-PAGE and immunoblotted with the indicated antibodies. Normal IgG was used as the negative control. **(A)** Co-immunoprecipitations of EloA, EloB with either PAF1 (left) or LEO1 (right) antibodies. **(B)** Co-immunoprecipitation of PAF1 with either EloA (left) or EloB (right) antibodies. **(C)** Nuclear extracts of HCT116 cells transfected for 48 h with plasmids expressing CUL5 and either PAF1 or LEO1 as indicated were immunoprecipitated using antibodies against PAF1 and LEO1. Immunoblotting was performed using the indicated antibodies. **(D, E)** HCT116 cells were treated with 2*μ*M doxorubicine (Dox) or mock-treated for 1 h **(D)** or 4 h **(E)**. Extracts were immunoprecipitated using antibodies against PAF1, LEO1 or an IgG control, as indicated. Immunoblotting was performed using antibodies to RNAPII, Elongin A and PAF1.

We next determined whether the interaction between PAF1C-EloA complex is modulated by DNA damage. Prolonged treatment of HCT116 cells with the DNA double strand break (DSB)-inducing agent Doxorubicin (Dox), induced loss of RNAPII which could be reversed by treatment with the proteasome inhibitor MG-132 (Figure S2B), indicating that RNAPII is degraded by the proteasome under these conditions. Cullin-Ring Ligase (CRL) complexes are activated by covalent attachment of the ubiquitin-like protein NEDD8 to the Cullin subunit. MLN4924 inhibits the NEDD8-activating enzyme, thereby blocking CRL activation (Soucy et al., 2009). We observed that RNAPII degradation was also abolished by MLN4924 (Figure S2C), confirming previous data suggesting that RNAPII degradation is triggered, at least in part, by a CRL complex. We next analysed the interaction between PAF1 and EloA in control cells and cells following short exposure (1 hour) to the DSB-inducing agent. Co-IP of either PAF1 or LEO1 following 1h of Dox treatment still detected RNAPII or EloA (Figure 2D), while these interactions were nearly abolished after 4 h of Dox, suggesting that PAF1-EloA interaction is dependent on RNAPII and/or transcription (Figure 2E).

### PAF1 facilitates RNAPII ubiquitination

Having demonstrated an interaction between PAF1 and the EloA ligase complex, we asked whether PAF1 affects RNAPII ubiquitination in cell extracts. To this end, cells were transfected with HA-tagged ubiquitin, EloA, CUL5 and Flag-RPB1 (Flag-RNAPII), in the presence or absence of PAF1, and in the absence of exogenous DNA damage. Cell extracts were prepared in 1% SDS-containing buffer to retain only covalent interactions. RNAPII was immunoprecipitated using the anti-FLAG antibody and analyzed by western-blotting using the anti-HA which detects the ubiquitinated form of the RNAPII. As observed in Figure 3A, a stronger signal appeared in the presence of PAF1 compared to the empty vector control (compare lane 4 to lane 3). No signal was observed in the absence of ubiquitin as expected. Overexpression of LEO, as for PAF1, led to an increase of RNAPII ubiquitination, suggesting that the PAF1C may facilitate the basal ubiquitination of RNAPII (Figure 3B). This result was confirmed by expressing HIS-tagged ubiquitin and purification of ubiquitinated RNAPII from extracts by denaturing nickel pulldown. Similarly, PAF1 overexpression led to increased RNAPII ubiquitination (Figure S3A).

**Figure 3.**
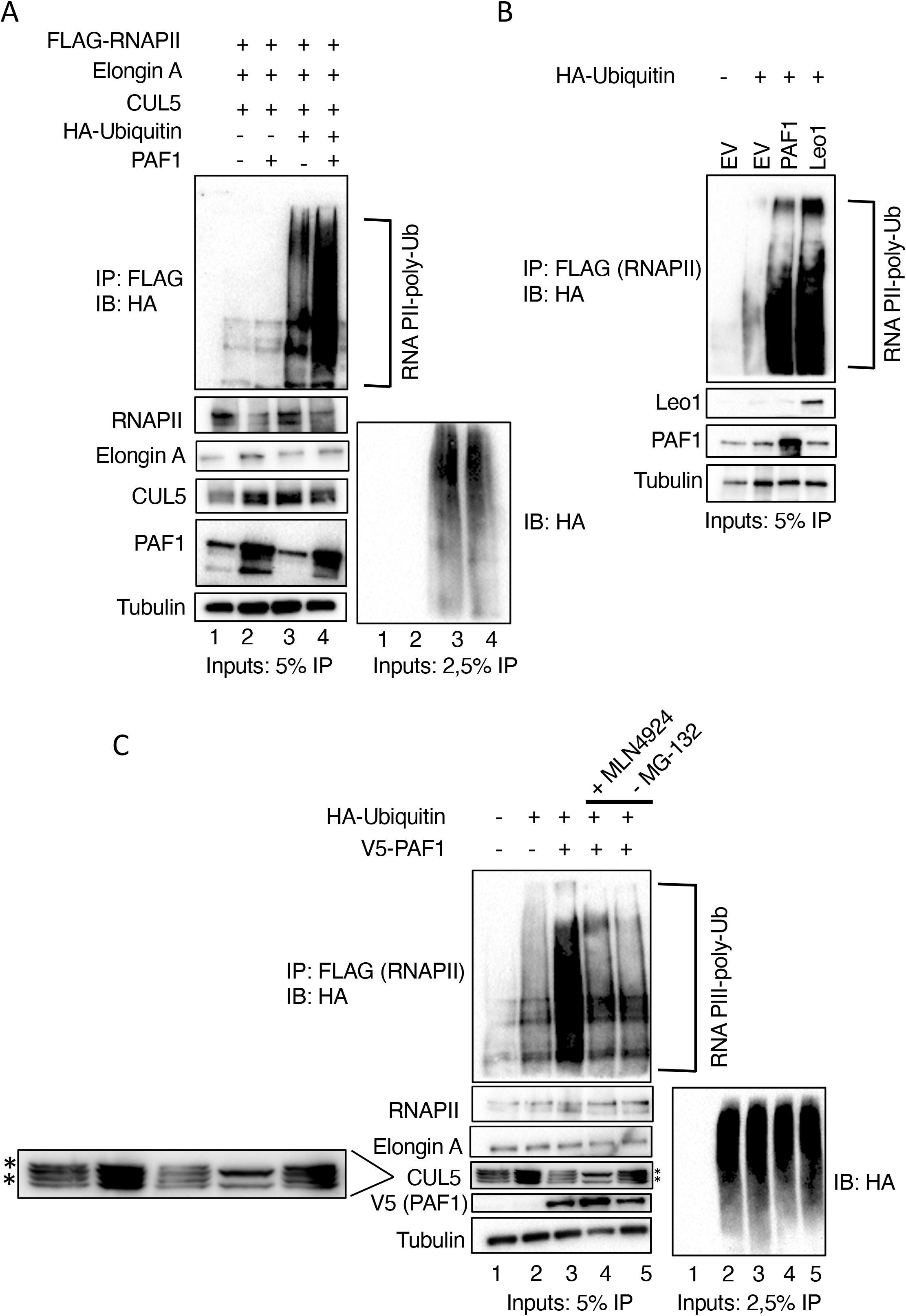
PAF1 facilitates RNAPII ubiquitination. **(A)** HEK293T cells were transiently transfected with EloA, CUL5, FLAG-RNAPII (all lanes), and HA-Ubiquitin and PAF1 plasmids as indicated for 48 h, treated with 40 *μ*M MG-132 for 4 h and harvested. Cell lysates were prepared as described in Material and Methods and immunoprecipitated with anti-FLAG antibody. Ubiquitinated forms of precipitated RNAPII were visualized by SDS-PAGE and immunoblotted using an anti-HA antibody. Western blotting of cell lysates (Inputs) performed with the indicated antibodies to detect the transfected proteins are shown below. Tubulin was included as a loading control. **(B)** Ubiquitination assay was performed as in (A) using either control (empty vector, EV) (lane 1, 2), PAF1 or LEO1 plasmids (lanes 3 and 4 respectively). Inputs are shown below. Tubulin was included as a loading control.**(C)** Ubiquitination assay was performed as in (A) using the indicated plasmids. Cells were treated for 16 h with 1 *μ*M MLN4924 (lane 4) or without MG-132 treatment (lane 5). Inputs are shown below. An enlargement of CUL5 blot is shown on the left. * CUL5 neddylated form.

Lysine linkages between ubiquitin monomers within polyubiquitin chains are associated with different physiological roles. K48-linked polyubiquitination leads to proteasomal degradation, whereas K63-linked polyubiquitination serves as a scaffold for signaling complex assembly, especially in DNA damage response pathways. We performed *in cellulo* ubiquitin assays using mutants of ubiquitin, mutated on all lysines except one, either K48 or K63. As observed in Figure S3B, only extracts transfected with ubiquitin carrying only the K48 presented the characteristic pattern of ubiquitinated RNAPII, in both control cells and those overexpressing PAF1. This result indicates that ubiquitination facilitated by PAF1-EloA tags RNAPII for proteasomal degradation.

As already observed, RNAPII ubiquitination depends on an activated CRL EloA complex since treatment with the inhibitor MLN4924, which inhibits the NEDD8-activating enzyme thereby blocking CRL activation (Soucy et al., 2009), led to an accumulation of RNAPII (Figure S2C). Furthermore, PAF1 overexpression strongly increased RNAPII ubiquitination (Figures 3A, S3A and S3B). In line with the finding that PAF1-induced RNAPII degradation occurs through the CRL EloA complex, we found that MLN4924, also inhibited RNAPII ubiquitination (Figure 3C, compare lanes 3 and 4). As observed in the corresponding input, the neddylated form of CUL5 is no longer observed either on the endogenous form, nor on the MYC-CUL5 overexpressed form. To confirm that CUL5 is indeed involved in RNAPII degradation, we constructed the dominant-negative mutant, CUL5ΔNEDD8 (K724A/K727A/K728A) which cannot be activated by NEDD8 (Ehrlich et al., 2009). Indeed, the ubiquitination signal showed a clear decrease (Figure S3C), and the corresponding input is not neddylated. In order to detect ubiquitinated proteins *in cellulo*, they must be allowed to accumulate by treating cells with proteasome inhibitor MG-132. Indeed, detection of ubiquitinated RNAPII in PAF1-expressing cells is significantly decreased in the absence of MG-132 (Figure 3C, compare lane 3 to lane5).

### PAF1 facilitates RNAPII ubiquitination through CTD phosphorylation by CDK12

CRLs often recognize phosphorylated substrates (Lydeard et al., 2013). Consistently, ubiquitination has been shown to occur on the hyperphosphorylated form of RPB1 involved in transcription elongation (Mitsui and Sharp, 1999). Consistent with this, proximity ligation assay (PLA) which allows the *in situ* detection of interacting endogenous proteins as well as post translational modifications (Soderberg et al., 2006), revealed a speckled pattern in U2OS cells labelled using both P-Ser2-RNAPII and ubiquitin antibodies, following transcriptional stress using a-amanitin a drug that induces RNAPII stalling and degradation (Weems et al., 2015) (Figures S4A and S4B), indicating that the Ser2 phosphorylated form of RNAPII is ubiquitinated. Prolonged a-amanitin treatment leads to complete degradation of RNAPII, which could be restored by treatment with the proteasome inhibitor, MG-132 (Figure S4A). Consistently, PLA signal was completely abolished in cells exposed to prolonged treatment with the drug, but was partially by treatment MG-132 (Figure S4B). A technical control in which each of the primary antibodies was substituted one at a time by an IgG control gave no signal, indicating that PLA signal obtained is specific (Figure S4B). These results confirm that ubiquinated RNAPII is likely phosphorylated on Ser2.

We then investigated which CTD kinase is required for RNAPII ubiquitination and degradation. RNAPII CTD is phosphorylated on Ser2 primarily by two kinases, CDK9 and CDK12. CDK9 is the kinase subunit of P-TEFb and is required for transcription elongation. CDK12 functions as a complex with cyclin K (CCNK) in the regulation of gene transcription (Blazek et al., 2011). Like CDK9, it phosphorylates RNAPII CTD at Ser2, which is thought to be a critical step in the transition from transcriptional initiation to elongation (Buratowski, 2009). PAF1C recruits CDK12 and regulates RNAPII pausing release (Yu et al., 2015). High levels of Ser2 phosphorylation by CDK12 are also required for subsequent mRNA 3’ end formation (Davidson et al., 2014). We therefore performed our ubiquitination assays by overexpressing either CDK9 or CDK12, compared to PAF1 (Figure 4A). As already observed, overexpression of PAF1 enhanced the ubiquitination of RNAPII (compare lanes 2 and 3). Overexpression of CDK9 did not change the RNAPII ubiquitination pattern compared to control cells (compare lanes 4 and 2). In contrast, CDK12 induced a very strong RNAPII ubiquitination signal (lane 5), stronger than that observed for PAF1 (lane 3) or in control cells (lane 2). Furthermore, overexpression of cyclin K also strongly enhanced RNAPII ubiquitination (Figure S4C), suggesting that CDK12/cyclin K complex facilitates RNAPII ubiquitination. CDK12-dependent RNAPII ubiquitination was mediated by CUL5-EloA, since it could be diminished by expression of the dominant-negative mutant CUL5ΔNEDD8 (Figure S4D). In addition, CTD-deleted RNAPII (Rosonina and Blencowe, 2004) was not targeted for CDK12-dependent ubiquitination (Figure 4B). Finally, addition of a CDK12 and CDK13 selective inhibitor, THZ531 (Zhang et al., 2016), significantly reduced PAF1-dependent RNAPII ubiquitination (Figure 4C). Immunoblotting of extracts confirmed that the Ser2 phosphorylation was strongly decreased by addition of THZ531 (Figure 4C). All together, these results show that RNAPII ubiquitination is mediated by the Elongin A ligase complex-PAF1-CDK12/CCNK pathway.

**Figure 4.**
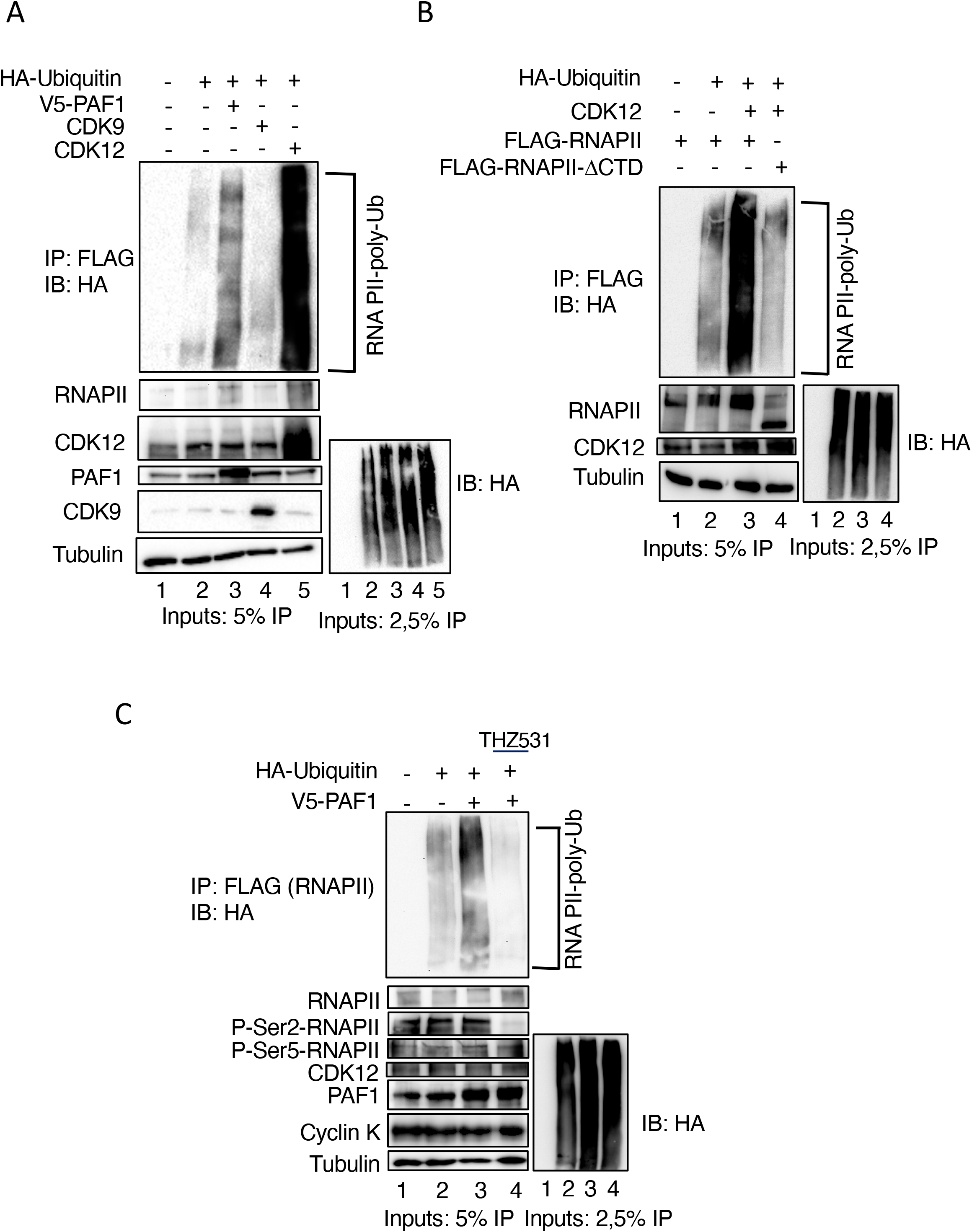
PAF1 facilitates RNAPII ubiquitination through CTD phosphorylation by CDK12. **(A)** HEK293T cells were transiently transfected with EloA, CUL5, FLAG-RNAPII (all lanes), and HA-Ubiquitin, PAF1, CDK9 or CDK12 plasmids as indicated, for 48 h. Cells were treated with 40 *μ*M MG-132 for 4 h and harvested. Cell lysates were prepared as described in Material and Methods and immunoprecipitated with anti-FLAG antibody. Ubiquitinated forms of precipitated RNAPII were visualized by SDS-PAGE and immunoblotted using an anti-HA antibody. Western blotting of cell lysates (Inputs) performed with the indicated antibodies to detect the transfected proteins are shown below. Tubulin was included as a loading control. **(B)** HEK293T cells were transiently transfected with plasmids expressing EloA, CUL5 (all lanes), HA-Ubiquitin, CDK12 and either full-length RNAPII or CTD-deleted RNA-PII as indicated, for 48 h. Ubiquitination assay was performed as in (A). Ubiquitinated forms of precipitated RNAPII were visualized by SDS-PAGE and immunoblotted using an anti-HA antibody. Western blotting of cell lysates (Inputs) performed with the indicated antibodies to detect the transfected proteins are shown below. Tubulin was included as a loading control. **(C)** HEK293T cells were transiently transfected with EloA, CUL5, FLAG-RNAPII (all lanes), and HA-Ubiquitin and CDK12 plasmids as indicated, for 48 h. Where indicated, THZ531 (1 *μ*M), the CDK12 inhibitor was added for 16h. Ubiquitination assay was performed as in (A). Ubiquitinated forms of precipitated RNAPII were visualized by SDS-PAGE and immunoblotted using an anti-HA antibody. Western blotting of cell lysates (Inputs) performed with the indicated antibodies to detect the transfected proteins are shown below. Tubulin was included as a loading control.

### CDK12 interacts with the Elongin A ligase complex

Interestingly, a recent proteomic analysis performed under stringent conditons identified EloA, EloB and EloC among the interactants of CDK12 (Bartkowiak and Greenleaf, 2015). In addition, CDK12 has been shown to interact with PAF1 (Yu et al., 2015). We therefore analysed interactions between CDK12 and PAF1, EloA and CUL5. As expected, CDK12 interacted with cyclin K, RNAPII, and PAF1. Notably, EloA and CUL5 were also co-immunoprecipitated by CDK12 (Figure 5A). CDK12, RNAPII, EloA and CUL5 were also co-immunoprecipitated with cyclin K (Figure 5B). Finally, the reverse IP using CUL5 antibody immunoprecipitated RNAPII and cyclin K (Figure 5C).

**Figure 5.**
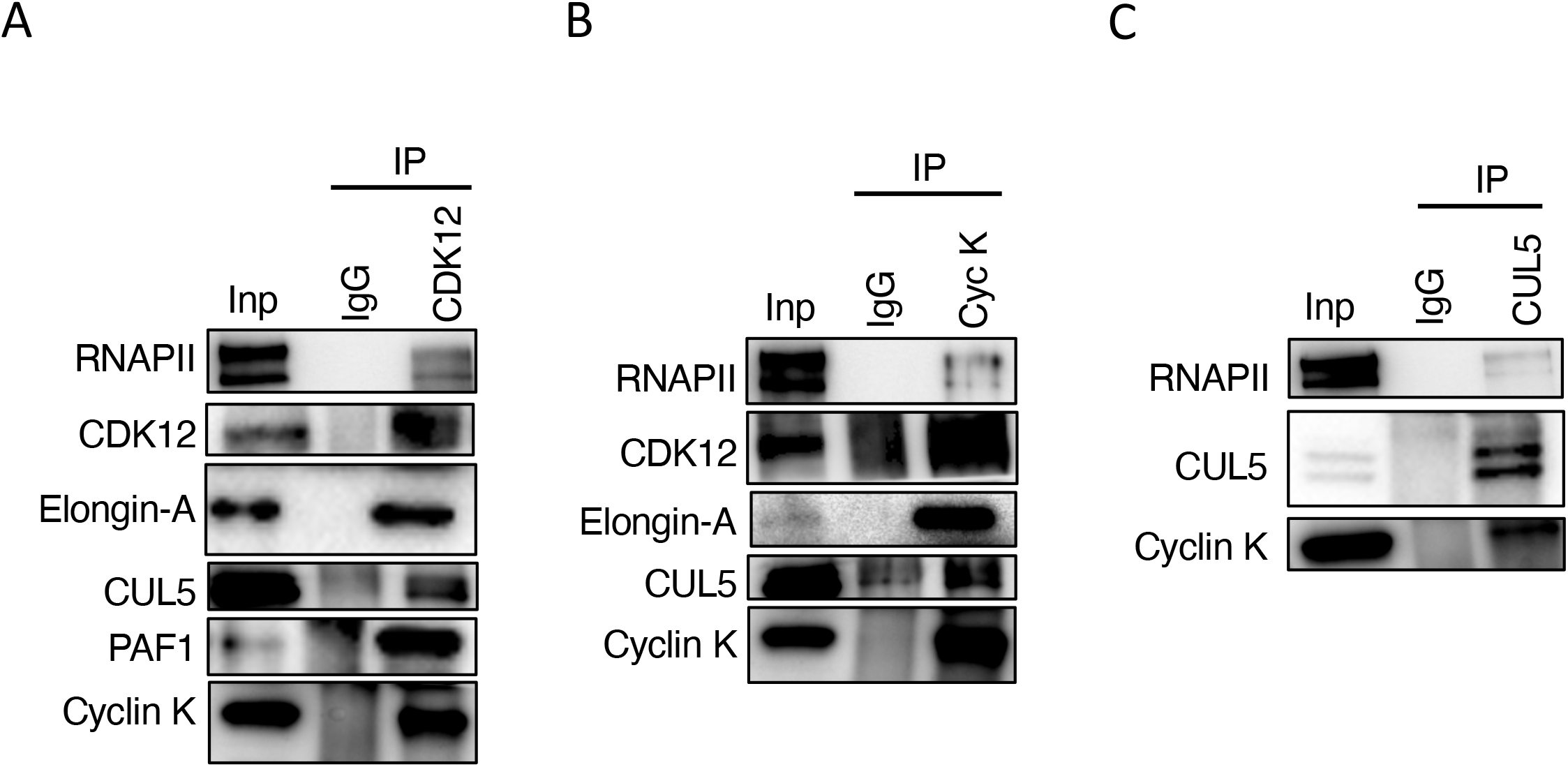
CDK12 interacts with the Elongin A complex. **(A-C)** HCT116 were transiently transfected with CDK12, EloA, CUL5 or Cyclin K as indicated for 48 h. Proteins from HCT116 nuclear extracts were immunoprecipitated with the indicated antibodies. Nuclear extracts (Inputs 5%) and immunoprecipitates were separated by SDS-PAGE and immunoblotted with the indicated antibodies. IgG were used as the negative control. **(A)** Co-immunoprecipitations of RNAPII, EloA, CUL5, XRN2, PAF1 and Cyclin K with CDK12 antibody. **(B)** Co-immunoprecipitations of RNAPII, CDK12, EloA, CUL5, XRN2 and PAF1 with Cyclin K antibody. **(C)** Co-immunoprecipitations of RNAPII, and Cyclin K with CUL5 antibody.

### P-Ser-2 RNAPII ubiquitination is decreased upon PAF1 depletion

To confirm previous data showing that PAF1 facilitates RNAPII degradation, we decided to analyse P-Ser2-RNAPII ubiquitination upon PAF1 depletion. Since RNAi approaches take days and may induce non-specific side effects, we chose to engineer conditional depletion of PAF1 on a faster time scale using gene editing. CRISPR/Cas9 was used to tag PAF1 at its N-terminus with an auxin-inducible degron (AID) (Figure 6A and Material and Methods for details). Upon exposure to hormones of the auxin family such as indole-3-acetic acid (IAA), AID-tagged proteins are degraded in a manner dependent on plant Tir1 protein (Natsume et al., 2016; Nishimura et al., 2009). Multiple resistant colonies were obtained and homozygous modification was shown by PCR on genomic DNA using primers designed outside of the homology arms. On all clones, it showed the two alleles tagged with hygromycin (upper band) and blasticidin (lower band) while the native cell line with untagged PAF1 (WT) presented a smaller product (Figure 6B). Another PCR using a primer inside of the tag mAID and a primer on the gene generated also a single band on the clones (Figure S5A). Western blotting of two clones confirmed homozygous targeting, shown by the higher molecular weight PAF1 and the absence of any signal at the size expected for the wild type PAF1 (Figure 5C). In cells treated with auxin, PAF1 protein was depleted very efficiently and specifically in cells expressing OsTIR1 (Figure 6C). We confirmed that these cells expressed mAID-PAF1 in the nucleus; a faint PAF1 expression or an antibody background is still visible under 2h of auxin treatment (Figure S5B). To test the mAID-PAF1 depletion, a time-course of auxin addition was performed on clone #2 and analyzed by western-blot. PAF1 levels are reduced within 30 min of IAA treatment and almost undetectable after 120 min (Figure S5C). The same samples blotted with CDC73, another component of the PAF1 complex did not show a depletion of the corresponding protein, indicating the specificity of the tagging (Figure S5C).

**Figure 6.**
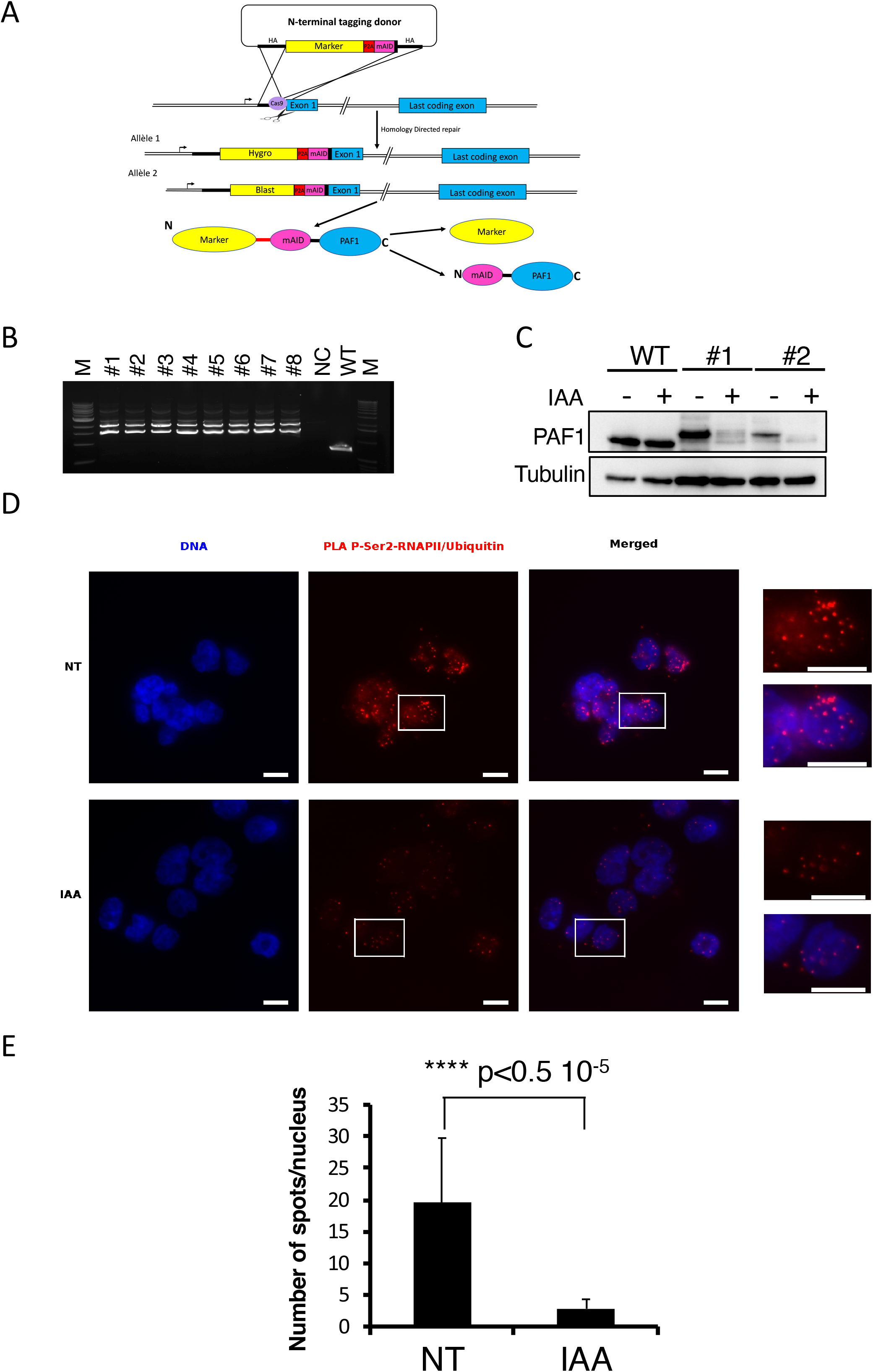
Depletion of PAF1 reduces ubiquitination of P-Ser2-RNAPII. **(A)** Schematic showing the CRISPR strategy for AID tagging of endogenous PAF1. Two repair cassettes containing the mAID tag, a P2A cleavage site, and either the hygromycin or blasticidin resistance markers were flanked by 5’ and 3’ homology arms (HA) for PAF1 gene. **(B)** Genotyping of N-terminally tagged alleles by PCR. The primer set amplifies a smaller PCR product (705 bp) from the WT allele, while it amplifies a larger product from hygromycin (upper band, 2321 bp) and blasticidin (lower band, 1704 bp) tagged alleles. NC, negative control with no DNA. M, marker. **(C)** Western-blotting of PAF1 in either parental Os-TIR1 HCT116 (WT) or in mAID-PAF1 cells from 2 different clones, using PAF1 antibody. The indicated cells were treated with or without 500 *μ*M IAA for 3 h. Tubulin was included as a loading control. **(D)** Representative image of the Proximity Ligation Assay (PLA) showing the P-Ser2-RNAPII/Ubiquitin complexes in mAID-PAF1 cells treated with or without 500 *μ*M IAA for 3 h. PLA signals are in red, DNA (blue) is dyed with DAPI. The merge between red and blue is shown. Right panels: enlarged views of the PLA signals. Scale bars, 10 *μ*M.**(E)** Quantification of PLA signals. Mean number of spots per nucleus from three independent experiments in non-treated (NT) and treated (IAA) cells. A minimum of 200 cells were counted for each experiment; error bars indicate SD ;(****, p <0.5 10^-5^, independent Student’s t test).

This cell line cannot be used in our ubiquitination assay since MG-132 treatment, required to observe the ubiquitination signal, would also inhibit the OsTIR1 E3 ligase complex. We therefore analyzed RNAPII ubiquitination by PLA on non-treated (NT) or IAA treated cells for 3 h. We detected specific PLA signals between P-Ser2-RNAPII and ubiquitin in the NT cells that were strongly decreased after IAA treatment (Figure 5D and 5E). The technical control in which each of the primary antibody was substituted one at a time by an IgG antibody was performed (Figure S5D). Unfortunately, EloA/CUL5 antibodies were not suitable for PLA. Collectively, these data indicate that PAF1 indirectly regulates RNAPII ubiquitination through Ser2 phosphorylation by CDK12.

## DISCUSSION

Each step in the RNAPII transcription cycle is largely regulated by accessory proteins that directly impact RNAPII activity, relay their response to other regulatory proteins or modify chromatin. Elongation is an extremely precisely controlled step at both early and productive stages (Kwak et al., 2013). Different protein complexes involved in the different stages of elongation have been widely described and studied. The Elongin complex is one of the factors required for more efficient elongation. The PAF1 complex is involved in the regulation of all stages of transcription, including elongation, and recently a number of studies have focused on its role in the release of the RNAPII proximal promoter pausing (Van Oss et al., 2017). However, PAF1C also regulates the progression of RNAPII by modulating the rate of elongation (Hou et al., 2019). In this paper, we show that these two complexes coordinately regulate RNAPII degradation and act as a mechanism of elongation surveillance in basal conditions through RNAPII CTD phosphorylation by CDK12.

Isolating the Elongin complex in the proteomic analysis of proteins interacting with PAF1 is consistent with previous reports showing that PAF1 interacts with positive elongation factors to promote transcriptional elongation (Hou et al., 2019; Rouillard et al., 2016). However, even if Elongin interacts directly with transcribing RNAPII and enhances the overall rate of transcript elongation by suppressing transient pausing (Conaway et al., 1993), it presents dual functions and acts also as the substrate recognition subunit of the CRL5 E3 ligase that targets RNAPII for degradation (for review, see (Conaway and Conaway, 2019) and references therein). Other E3 ligases have been described for RNAPII ubiquitination and degradation such as BRCA1/BARD1 (Kleiman et al., 2005) or WWP2 (Caron et al., 2019), but in all cases, studies were done on cells under genomic or transcriptional stress (Caron et al., 2019; Gregersen and Svejstrup, 2018; Nakazawa et al., 2020; Tufegdzic Vidakovic et al., 2019). Using *in cellulo* ubiquitin assays we show that PAF1 facilitates RNAPII ubiquitination, under basal conditions through the CRL5-Elongin complex since CUL5 mutation decreases this RNAPII ubiquitination.

PAF1C acts as a platform recruiting factors that connect PAF1 to multiple transcription processes from elongation to polyadenylation and termination. During productive elongation, both the PAF1 and Elongin complexes are associated with chromatin. In response to DNA damage the protein Cockaine Syndrome B (CSB) regulates the recruitment of EloA and the ligase complex at the site of DNA damage but is not essential for the assembly of the EloABC-CUL5 ligase complex (Harreman et al., 2009). During the course of this work, a new transcription recovery pathway was reported that involves the CSB-dependent association of the PAF1C with RNAPII after UV irradiation, required to restore a processive transcription after genotoxic stress (bioRxiv preprint: https://doi.org/10.1101/2020.01.04.894808), linking PAF1 and Elongin pathways through CSB after DNA damage. We show here that, in the absence of induced DNA damage, there is interaction between these two complexes including the E3 ligase form suggesting that transcription *per se* is sufficient to induce the assembly of the ligase complex. Consistent with this notion, we recently showed that the DNA repair complex, MRE11-NBS1-Rad50 (MRN), is associated with chromatin in a transcription-dependent manner and protects highly transcribing genes from genomic instability (bioRxiv preprint: https://www.biorxiv.org/content/10.1101/2020.05.27.118638v1). Thus, during transcription, should RNAPII become stalled and require degradation, the PAF1-Elongin ligase complex would ensure that RNAPII would be targeted for degradation by ubiquitination and rapidly dissociated from the chromatin. For example, in yeast, the ATP-dependent INO80 complex and CDC48/p97 remodeling complex are required for turnover of chromatin-bound RNAPII to prevent the accumulation of ubiquitinated RPB1 on chromatin (Lafon et al., 2015). Moreover, still in yeast, when replication forks encounter transcription, the INO80 with CDC48 promotes removal of RNAPII from the chromatin with the help of its association with PAF1. Both INO80 and PAF1 are needed to achieve efficient removal of the transcription complex and its transient degradation (Poli et al., 2016).

It is noteworthy that PAF1C actively orchestrates multiple histone-modifying factors in the formation of histone marks linked to transcription (Kim et al., 2009) including H2B monoubiquitination (Ng et al., 2003) that is associated with regions of active transcription. Moreover, H1.2 localizes at target genes through its interaction with phosphorylated Ser2-RNAPII (P-Ser2-RNAPII) and stably interacts with Cul4A E3 ubiquitin ligase and PAF1C to potentiate target gene transcription by facilitating H4K31 ubiquitination (Kim et al., 2013). This study shows that PAF1 is also able to facilitate ubiquitination modification of P-Ser2-RNAPII directly to negatively regulate transcription elongation.

P-Ser2-RNAPII-CTD is strongly associated with transcription elongation and it has been proven, at least in yeast, that only the Ser2 phosphorylated elongating form of RNAPII is the target of efficient Rpb1 ubiquitination (Mitsui and Sharp, 1999; Somesh et al., 2005). Much attention has been given to the importance of promoter-proximal pausing of RNAPII in regulating transcription elongation. P-TEFb (CDK9/Cyclin T) dependent phosphorylation events release paused Pol II, and create a platform for binding of RNA processing factors on the P-Ser2 CTD of Pol II (Adelman and Lis, 2012). However, other studies indicate that CDK12 is the major P-Ser2 kinase (Bartkowiak et al., 2010). Apparently contradictory studies involve PAF1 either in maintaining the pause (Chen et al., 2015) or in promoting the pause release (Yu et al., 2015), underlining the complexity of PAF1’s role in transcriptional regulation. In the case of pause release, PAF1C recruits CDK12 promoting transcript elongation. Another report indicates that Ser2 phosphorylation occurs after pause release (Lu et al., 2016), suggesting that PAF1 function in pause release may relate more to its interaction with P-TEFb than to its implication in CDK12 recruitment. Here, we show that phosphorylation by CDK12 contributes very strongly and possibly much more directly than PAF1 to the ubiquitination of RNAPII. Moreover, P-Ser2 also governs the recruitment of processing factors at polyadenylation (pA) sites (Ahn et al., 2004). The 3’ end formation of pre-mRNA and P-Ser2 by CDK12 are thus reciprocally coupled (Davidson et al., 2014). In addition, PAF1C interacts with cleavage and pA factors (Rozenblatt-Rosen et al., 2009) and stimulates cleavage and pA transcript (Nagaike et al., 2011). Thus, the recruitment of CDK12 by PAF1 could maintain RNAPII in a hyperphosphorylated form, and, in the event of transcription stalling, to trigger an early termination by causing the degradation of RNAPII with the Elongin A ligase complex.

In this study, we identified a mechanism by which PAF1C and CDK12 coordinate with Elongin A to generate a surveillance mechanism network during transcription elongation that will rapidly degrade RNAPII in case of transcription arrest. Whether and how this mechanism applies to signal-specific or gene-specific stimulation requires future investigation.

## MATERIAL AND METHODS

### Antibodies and Plasmids

Antibodies used in this study are listed in Table S4.

pcDNA3 PAF1 and pcDNA3 LEO1 plasmids were gifts from Matthew Meyerson (Addgene plasmids # 11061 and 11059, respectively). FLAG-Pol-II WT and FLAG-Pol-II-Δ (entire CTD deleted) were gifts from Benjamin Blencowe (Addgene plasmids # 35175 and # 35176 respectively). pcDNA3-MYC-CUL5 was a gift from Yue Xiong (Addgene plasmid # 19895). TCEB3 (Elongin A) and CCNK (Cyclin K) plasmids were obtained from MGC (Dharmacon, MHS6278-202830313 and MHS6278-202756944, respectively). CDK12 plasmid was kindly given by Dalibor Blazek (Genes and Dev, 2011) to RK. 8xHA-ubiquitin (pMT123) and 8xHIS-ubiquitin (pMT107) were kindly provided to SR by Dirk Bohmann (Treier et al., 1994).

V5-PAF1 plasmid was obtained by cloning PAF1 in Gateway pcDNA3.1/nV5-DEST vector (Invitrogen). The untagged CDK9 plasmid was obtained by cloning the cDNA of HA-CDK9 (gift from Andrew Rice, Addgene #28102) in pCDNA3.1. Myc-CUL5-ΔNEDD8 (K724A/K727A/K728A) mutant was obtained using QuickChange Multi Site-Directed Mutagenesis kit from Agilent (#200514) using the primer shown in Table S5. espCas9(1.1) was a gift from Feng Zhang (Addgene plasmid # 71814). pMK344 and pMK347 (Addgene plasmids #121118 and #121181) were gifts from Masato Kanemaki (Yesbolatova et al., 2019).

### Chemicals and Reagents

Cycloheximide, Doxorubicine, MG132, and IAA were purchased from Sigma-Aldrich. THZ531 was obtained from MedChemExpress and MLN4924 from Tebu-bio.

### Cell Culture and Transfection

HeLa S3, U20S, HEK293T and HCT116 were obtained from the ATCC; HCT116-OsTIR1 was kindly provided by Masato Kanemaki (Natsume et al., 2016). Cells were routinely grown in a humidified incubator at 37°C in a 5% CO2 atmosphere. HeLa S3, U2OS and HEK293T cells were propagated in Dulbecco’s modified Eagle’s medium (DMEM) supplemented with 10% FBS and antibiotics. HCT116 cells were grown in McCoy 5A supplemented with 10% FBS and antibiotics. Plasmid transfection was performed with Jet PEI (PolyPlus Transfection) or Fugene HD (Promega) according to the manufacturer’s instructions. All samples were harvested at approximately 48 h (plasmids).

### Protein complex purification

PAF1 complex was purified from Dignam high salt nuclear extracts (dx.doi.org/10.17504/protocols.io.kh2ct8e) from HeLa-S3 cells stably expressing FLAG-HA-PAF1 by two-step affinity chromatography (dx.doi.org/10.17504/protocols.io.kgrctv6). Sequential FLAG and HA immunoprecipitations were performed on equal amounts of proteins. Silver-staining was performed according to the manufacturer’s instruction (Silverquest, Invitrogen). Mass spectrometry was performed at Taplin facility, Harvard University, Boston. Protein functions were obtained using the Gene Ontology (GO) database of biological processes. Processes represented by at least three different proteins were presented in the form of a pie chart, where the size of each slice is proportional to the number of proteins associated with that biological process. These results are also presented in tabular format, listing these processes and their associated proteins explicitly. In parallel, pathways analysis was performed using KEGG (Kyoto Encyclopedia of Genes and Genomes) and Reactome (reactome.org) databases and were represented in the form of a pie chart or in tabular format as indicated above.

### Immunoprecipitation

Immunoprecipitations were performed on nuclear extracts on approximately 400 *μ*g proteins. Cells were lysed in ice-cold hypotonic buffer (20 mM TRIS pH 7.6, 10 mM KCl, 1,5 mM MgCl2) supplemented with EDTA-free complete protease inhibitor mixture (PIC, Roche) and phosphatases inhibitors (PhosphoStop, Roche) for 15 min on ice. NP40 was added at 0.5% final and extracts were centrifuged 1 min at 14,000g/4°C. The pellet (nuclei) was resuspend in nuclease buffer (20 mM TRIS pH 7.6, 150 mM NaCl, 1,5 mM MgCl2, 2.5 mM CaCl2, 0,5 *μ*l PMSF 100 mM) and incubated with Micrococcal nuclease (New England Biolabs) or Benzonase (Sigma-Aldrich) 2h at 4°C. Lysates were cleared by centrifugation at 14,000g/4°C for 10 min and diluted in IP buffer (50 mM TRIS pH 7.6, 150 mM NaCl, 1% NP40) supplemented protease and phosphatase inhibitors. Protein concentration was determined using the Bradford reagent (Biorad). Immunoprecipitations were performed on 400 *μ*g of protein extracts with the indicated antibodies and rotated overnight at 4°C. Protein-G Dynabeads were washed three times in IP buffer, added to protein extracts/antibody solution and incubated for 2 h at 4°C. Immunoprecipitates were washed extensively with the IP buffer, resuspended in protein sample loading buffer, boiled for 5 min, and analyzed by Western blotting.

### Immunoblotting

For immunoblot, protein extracts were either nuclear extracts (see above) or whole-cell extracts obtained using RIPA buffer (50 mM TRIS pH 7.5, 150 mM NaCl, 1 % NP40, 0.5 % Sodium Deoxycholate, 0.1 % SDS) supplemented with Complete Protease Inhibitor (Roche). Protein concentration was determined using a Bradford assay. Protein extracts were resolved using SDS-PAGE, transferred to PVDF membrane, and probed using the indicated primary antibodies (Table SX) and anti-mouse or anti-rabbit IgG-linked HRP secondary antibodies (Southern Biotech) followed by ECL (Luminata Crescendo or Forte, Millipore).

### *In vivo* ubiquitination assays

Subconfluent HEK293T cells in 100 mm plates were transfected with 1.5 *μ*g 8xHA-Ubiquitin, 2 *μ*g FLAG-RNA-Polymerase II, 1 *μ*g elongin A, 1 *μ*g MYC-Cullin-5, and 5 *μ*g of the indicated plasmid (PAF1, CDK12, CDK9, empty vector). For each transfection, the total plasmid was adjusted to 10.5 *μ*g/100 mm by adding the parental pCDNA3.1 vector (empty vector, EV). 48 h after transfection, cells were treated with 40 *μ*M MG132 for 4h then lysed in 50 mM TRIS pH 7.4, 120 mM NaCl, 5 mM EDTA, 0.5% NP40, 0.5% Triton-X100, 1% SDS and protease and phosphatase inhibitors. Protein concentration was determined using the Bradford Ultra reagent (Expedeon). Immunoprecipitations were performed on 400 *μ*g of protein extracts using anti-FLAG antibody and the precipitated ubiquitinated forms were visualized using an anti-HA antibody and western blot analysis. Note that the levels of ubiquitination were assessed on equal amounts of proteins, as shown by the western blots of an unpurified fraction of the same extracts using anti-a-tubulin antibodies (Inputs).

### Purification of HIS-tagged ubiquitinated proteins

Subconfluent HEK293T cells in 100 mm plates were transfected with 5 *μ*g 8xHIS-Ubiquitin, 2 *μ*g FLAG-RNA-Polymerase II, 1 *μ*g Elongin A, 1 *μ*g MYC-Cullin-5, and 5 *μ*g of the indicated plasmid. 48h after transfection, cells were treated with 40 *μ*M MG132 for 4h then lysed under denaturing conditions (6M guanidinium-HCl, 10 mM TRIS pH8.0, 100mM sodium phosphate buffer pH8.0). HIS-Ub-conjugated proteins were purified by cobalt chromatography (TALON metal affinity resin, Clontech), as described in the manual. Cobalt-bound ubiquitinated proteins were subjected to western blot analysis using anti-RNA Pol II antibodies. Note that the levels of ubiquitination were assessed on equal amounts of proteins, as shown by the western blots of (1) the cobalt-bound ubiquitinated proteins using anti-HIS antibodies and (2) an unpurified fraction of the same extracts using anti-a-tubulin antibodies (Inputs).

### Immunofluorescence

Cells grown in coverslips were fixed with 4% PFA for 15 min, washed in PBS, and permeabilized in PBS-0.5% Triton X-100. The slides were then saturated with PBS, 3% BSA, 0,05% Tween20 containing the primary antibodies in a humid chamber at 4°C overnight and washed three times with PBS. Secondary antibodies (Alexa Fluor 488, 594-conjugated goat anti-rabbit or anti-mouse immunoglobulins G (IgGs) (1/1,000; Invitrogen) at 37°C for 30 min. Coverslips were mounted using ProLong Gold (Invitrogen). IF signals were acquired via a Zeiss CDD Axiocam Mrm monochrome from a Zeiss Axioimager Z1 microscope using a 63x Plan Apochromat 1.4 NA oil objective lens and Zen software. Exposure times, brightness and contrast settings are identical between images. Image mounting was done using OMERO (University of Dundee and Open Microscopy Environment).

### Generation of HCT116 mAID-PAF1 cell line

PAF1 was homozygously N-terminally tagged at its endogenous locus with the auxin-inducible degron (mAID) in OsTIR1-expressing HCT116 cell line as described (Yesbolatova et al., 2019). Specific guide RNA (sgRNA) targeting the region adjacent to the 5’ end of PAF1 gene (5’-GCGCCATAGCGACGAGGCGACGG-3’) and Cas9 were expressed from espCas9(1.1). Hygromycin or blasticidin resistance markers were incorporated into the cassette for homology directed repair (HDR) so that bi-allelic modification could be selected for (Yesbolatova et al., 2019). A P2A site, between the AID and drug markers, ensured their separation *via* peptide cleavage during translation (Kim et al., 2011) (Figure 6A). Donor plasmids carrying ~500 bp homology arms cloned by PCR from HCT-OsTIR1genomic DNA, mini-AID, and either Hygromycin or Blasticidin marker cassettes were created by inverse PCR using pMK347 and pMK344 as described (Yesbolatova et al., 2019). Primers used for the constructs are listed in Table S5. HCT-OsTIR1 cells were co-transfected with these two constructs together with the PAF1-sgRNA expressing Cas9 plasmid using Fugene HD (Promega), and selected with 100 *μ*g/ml Hygromycin B (Invivogen) and 10 *μ*g/ml Blasticidin (Invivogen). Single clones resistant to both drugs were then isolated and analyzed by PCR on genomic DNA and immunoblotting for PAF1 to confirm homozygous AID tagging. 500 *μ*M indole-3-acetic acid was added to cells to induce degradation of AID-PAF1.

### Proximity Ligation Assays (PLAs)

Cells grown in coverslips were fixed with 4% PFA for 15 min, washed in PBS, and permeabilized in PBS-0.5% Triton X-100. The slides were then saturated with PBS, 3% BSA, 5 *μ*g/ml salmon sperm DNA, 1 mM EDTA and probed with the indicated primary antibodies diluted in PBS, 1% BSA, 0,05% Tween20, 2.5 *μ*g/ml salmon sperm DNA, 1 mM EDTA. The PLAs were conducted using the Duolink In Situ kit (DUO92008, Sigma-Aldrich) and secondary antibodies coupled to DNA probes (DNA probe rabbit DUO92002; DNA probe mouse DUO92004) as indicated by the manufacturer. Coverslips were mounted using Prolong Gold (Invitrogen). Z-stacks images of PLA signals were acquired with a 63x Plan Apochromat 1.4 NA oil objective lens and Zen software. Image mounting was done using OMERO. For quantitative analyzes, Z-stacks were acquired every 0.24 *μ*m (Z step) with a range of 7 *μ*m. The number of PLA dots were quantified with Imaris software (Bitplane, South Windsor, CT, USA) that allows detection of objects in 3-dimensional space, based on size Surface reconstruction tool (DAPI channel) and new Spots tool (Texas Red channel) *via* an appropriate thresholding. For each condition (non-treated versus IAA treated cells), all the individual signals of a nucleus were counted for each nucleus. The data were exported to an Excel table and statistical analyses were performed using Excel functions. p value was calculated using the Student’s T test of the Excel software.

## Acknowledgments

We thank Dr Masato Kanemaki for providing the HCT116-OsTIR1 cell line and helpful advices; Dr Dalibor Blazek for CDK12 plasmid and Dr Bohman for HA- and HIS-tagged ubiquitin plasmids; the Montpellier Ressources Imagerie plateforme (MRI), member of the national infrastructure France-BioImaging supported by the French National Agency (ANR-10-INSB-04, Investments for the future). SR thanks Amélie Sarrazin from MRI for her precious help in microscopy and image analysis, Maria Morel-Carretero for PLA advices and Dominique Giorgi for statistical analyzes of PLA experiments and for his constant support during these past years.

## Author contribution

GS performed and analyzed experiments. JB performed experiments. CE analyzed MS data. RK initiated the project and wrote the paper; funding acquisition. SR designed, performed and analyzed experiments; wrote the paper (original draft).

