## Supporting Information for "PAF1 facilitates RNA polymerase II ubiquitination by the Elongin A complex through phosphorylation by CDK12"

**Table S1. List of interactants of Flag-HA-PAF1 identified by mass spectrometry.**

**Table S2. Biological processes and list of interactants of Flag-HA-PAF1 identified by mass spectrometry.**

**Table S3. Pathways and list of interactants of Flag-HA-PAF1 identified by mass spectrometry.**

**Table S4. List of antibodies used in this study.**

**Table S5. Sequences of oligos used in this study**

**Figure S1. Identification of PAF1 partners.**

**(A)** Left panel: western blot analysis of HeLa S3 mock cells or stably expressing FLAG-HA-PAF1 (FH-PAF1) using PAF1 antibody. Right panel: experimental scheme. FH-PAF1 was sequentially purified with anti-FLAG and anti-HA antibodies from Dignam nuclear extracts prepared from HeLa S3 mock cells or stably expressing FH-PAF1. Cellular nuclear partners of FH-PAF1 were determined by tandem mass spectrometry.**(B)** Pie chart of pathways (KEGG, Kyoto Encyclopedia of Genes and Genomes; Reactome, reactome.org) of proteins identified by mass spectrometry. The proteins involved are listed in Table S3.

**Figure S2. Degradation of RNAPII under Dox treatment is Cullin-Ring-Ligase (CRL)-dependent.**

**(A)** HCT116 cells were transfected with EloA plasmid for 48 h. Nuclear extracts (Inputs 5%) and EloA immunoprecipitates were separated by SDS-PAGE and immunoblotted with PAF1 antibody.**(B)** HCT116 cells were treated with 2$\mu$M doxorubicine (Dox) for 4 h in the presence of the protein synthesis inhibitor cycloheximide (CHX, 25 $\mu$g/ml) with or without the proteasome inhibitor MG132 (10 $\mu$M). Total cell lysates were prepared, separated by SDS-PAGE and immunoblotted with RNAPII (F-12), PAF1 and tubulin (loading control) antibodies. **(C)** Left panel: HCT116 cells were pretreated or not for 16 h with the NEDD8 activating enzyme (NAE) inhibitor MLN4924 (1 $\mu$M) and treated with Dox for 4 h in the presence of CHX as indicated in (B). Total cell lysates were prepared, separated by SDS-PAGE and immunoblotted with RNAPII (F-12), CUL5 and tubulin (loading control) antibodies. Right panel: scheme of the poly-ubiquitination of RNAPII by the Elongin A ligase complex. RNAPII is first monoubiquitinated by NEDD4 (N4) before being polyubiquitinated by the CRL5 complex ligase. NEDD8 (N8) needs to be activated by the NAE to be able to activate CUL5.

**Figure S3. PAF1 facilitates RNAPII poly-ubiquitination.**

**(A)** HEK293T cells were transiently transfected with EloA, CUL5, FLAG-RNAPII and 6xHis-Ubiquitin (all lanes) and either empty vector or PAF1 plasmids for 24 h. Cells were treated with 40 $\mu$M MG132 for 4 h. Ni+-purified 6xHis-Ub-conjugated proteins were probed either for the presence of ubiquitinated RNAPII using an anti-RNAPII (F-12) (left) or an anti-His antibody (right). Cell lysates were analyzed by western blotting with the indicated antibodies (Inputs). **(B)** HEK293T cells were transiently transfected with EloA, FLAG-RNAPII (all lanes), empty vector or PAF1 plasmids and either the K48-Ubiquitin or the K63-Ubiquitin as indicated for 48~ h. Cells were treated with 40 $\mu$M MG-132 for 4 h and harvested. Cell lysates were prepared as described in Material and Methods and immunoprecipitated with anti-FLAG antibody. Ubiquitinated forms of precipitated RNAPII and cell lysates (Inputs) were visualized by SDS-PAGE and immunoblotted using an anti-HA antibody. **(C)** HEK293T cells were transiently transfected with EloA, FLAG-RNAPII (all lanes), HA-Ubiquitin and PAF1 as indicated, and either CUL5-WT (lanes 1 to 3) or CUL5-MUT (lane 4) plasmids for 48 h, treated with 40 $\mu$M MG-132 for 4 h and harvested. Cell lysates were prepared as described in Material and Methods and immunoprecipitated with anti-FLAG antibody. Ubiquitinated forms of precipitated RNAPII were visualized by SDS-PAGE and immunoblotted using an anti-HA antibody. Western blotting of cell lysates (Inputs) performed with the indicated antibodies to detect the transfected proteins are shown below. Tubulin was included as a loading control. * CUL5 neddylated form.

**Figure S4. PAF1 facilitates RNAPII ubiquitination through Ser2 CTD phosphorylation by CDK12.**

**(A)** Upper blot: protein extracts from U2OS treated for 16 h with increasing doses of α-amanitin as indicated, were analyzed by immunoblotting using RNAPII and tubulin (loading control) antibodies. Lower blot: protein extracts from U2OS treated for 16 h with 15 $\mu$g/ml of α-amanitin without or with MG-132 (10 $\mu$M), were analyzed by immunoblotting using RNAPII and tubulin (loading control) antibodies. **(B)** U2OS cells were treated with 15 $\mu$g/ml of α-amanitin as indicated, without or with MG-132. PLA was performed using P-Ser2-RNAPII and ubiquitin combination antibodies (red signal). DNA was labelled with Hoechst (blue). The merge is shown at the right. Controls with each antibody alone in combination with IgG is shown below. Scale bar, 10 $\mu$M. **(C)** HEK293T cells were transiently transfected with EloA, CUL5, FLAG-RNAPII, HA-Ubiquitin (all lanes), and cyclin K plasmids as indicated, for 48 h. Ubiquitination assay was performed as in Figure 4. Ubiquitinated forms of precipitated RNAPII were visualized by SDS-PAGE and immunoblotted using an anti-HA antibody. Western blotting of cell lysates (Inputs) performed with the indicated antibodies to detect the transfected proteins are shown below. Tubulin was included as a loading control.**(D)** HEK293T cells were transiently transfected with EloA, FLAG-RNAPII (all lanes), HA-Ubiquitin and CDK12 as indicated, and either CUL5-WT (lanes 1 to 3) or CUL5-MUT (lane 4) plasmids for 48 h. Ubiquitination assay was performed as in Figure 4. Ubiquitinated forms of precipitated RNAPII were visualized by SDS-PAGE and immunoblotted using an anti-HA antibody. Western blotting of cell lysates (Inputs) performed with the indicated antibodies to detect the transfected proteins are shown below. Tubulin was included as a loading control.

**Figure S5. mAID-PAF1 cell line characterization.**
**(A)** Genotyping of N-terminally tagged alleles by PCR. The primer set generates a PCR product from the tagged alleles exclusively. **(B)** Immunofluorescence (IF) analysis of mAID-PAF1 cells (clone #2) treated with or without 500 $\mu$M IAA for 3 h, using the anti-PAF1 antibody (red). DNA (blue) is dyed with DAPI. Scale bar, 10 $\mu$M. **(C)** Time course of 500 $\mu$M IAA addition on mAID-PAF1 cells (clone #2). mAID-PAF1 was detected by anti-PAF. Tubulin was included as a loading control. The arrow indicates the PAF1 protein; *, aspecific band. **(D)** Technical control for PLA signals (red) shown in Figure 5D. Each antibody alone in combination with IgG is shown. DNA (blue) is dyed with DAPI. Scale bar, 10 $\mu$M.
