## Supplementary figures and images for "PAF1 facilitates RNA polymerase II ubiquitination by the Elongin A complex through phosphorylation by CDK12"

### Supplemental Figures

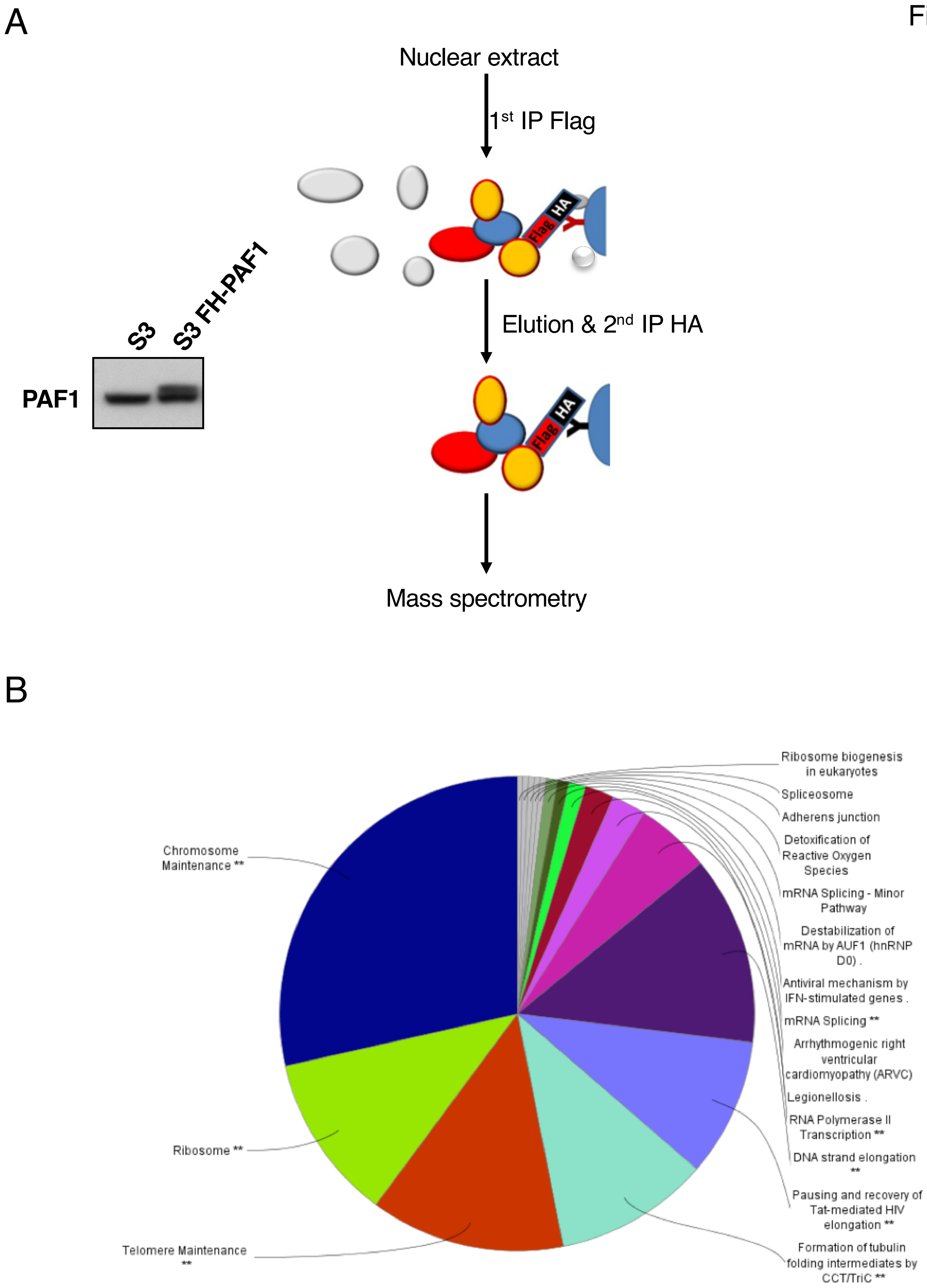

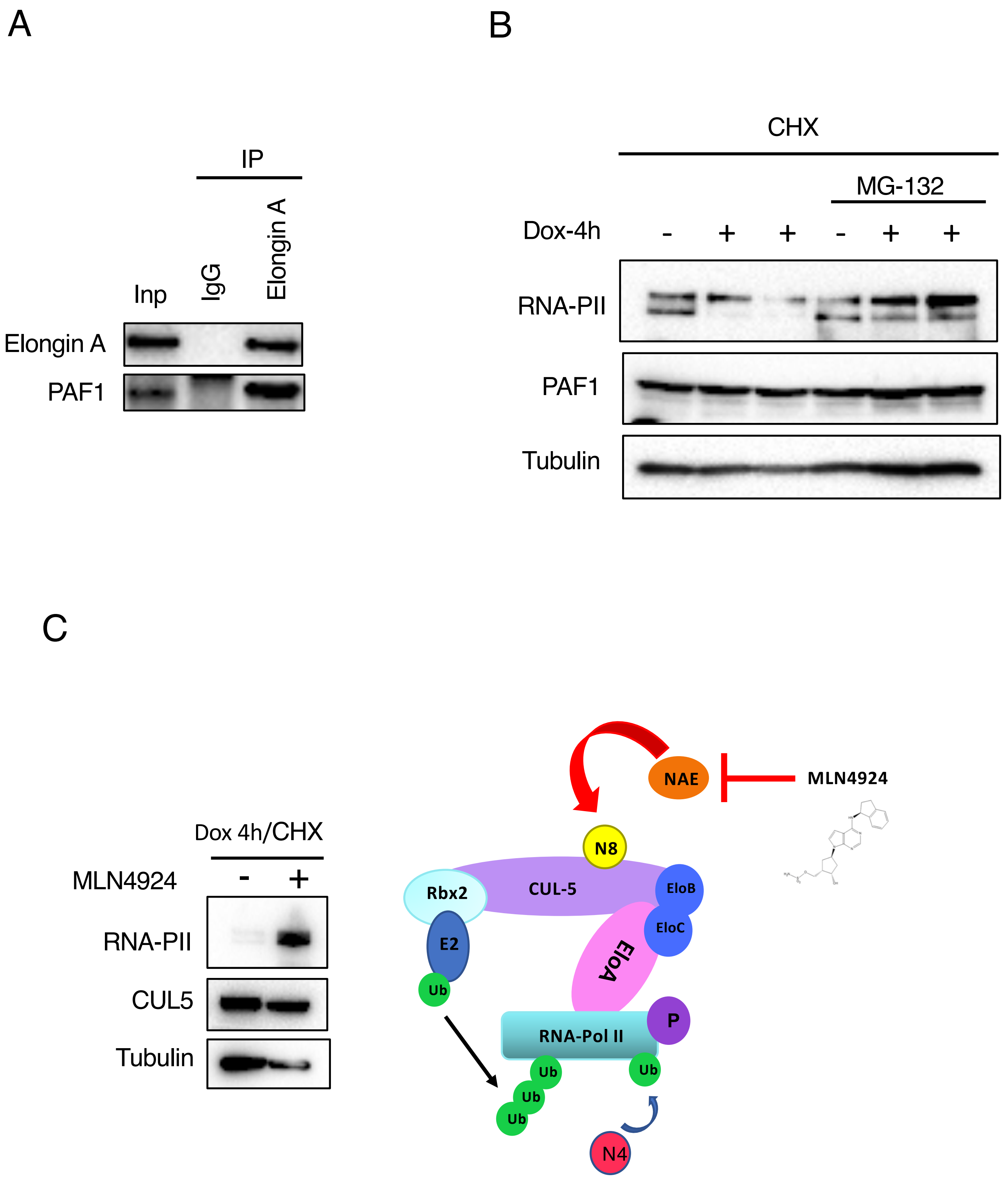

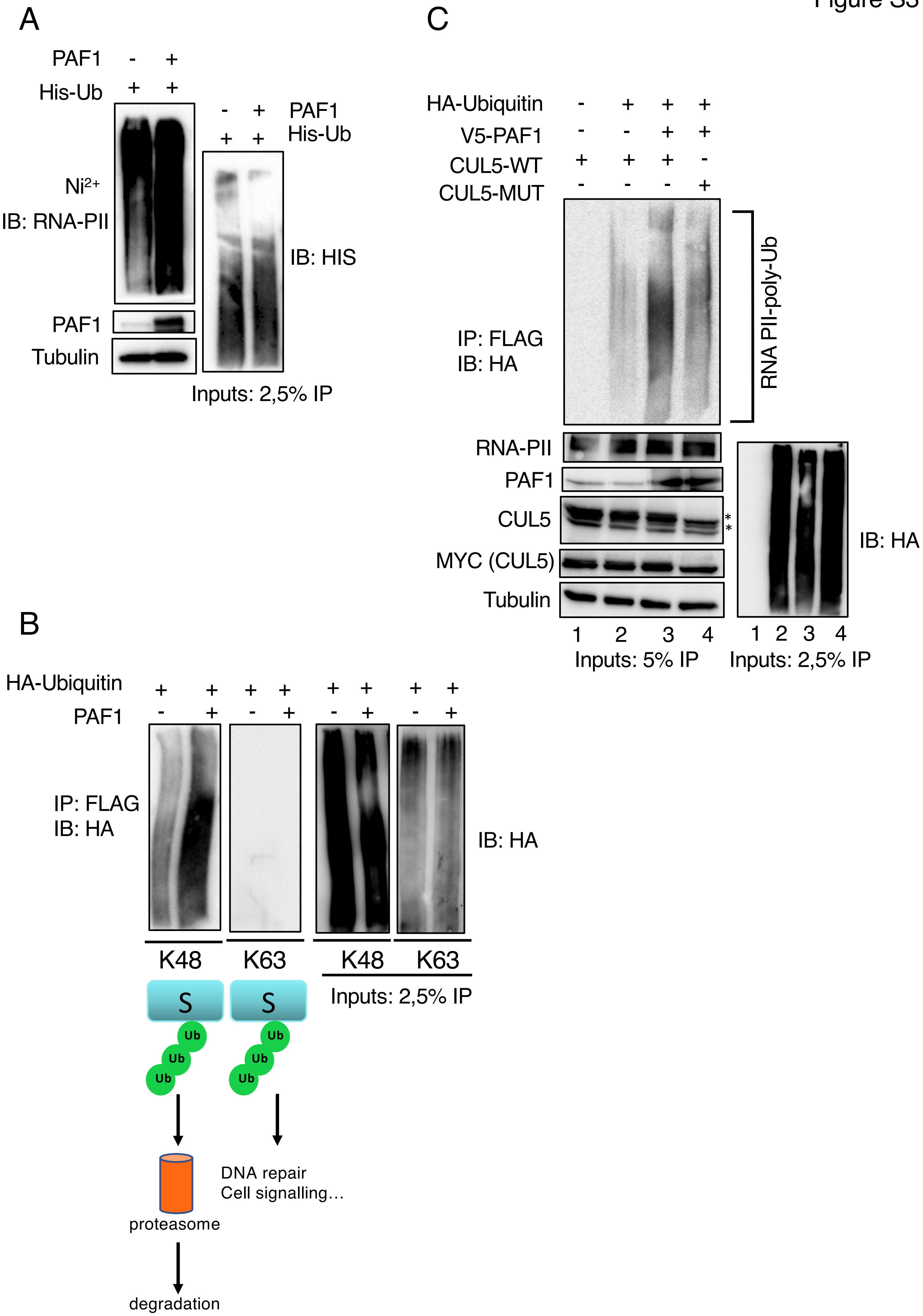

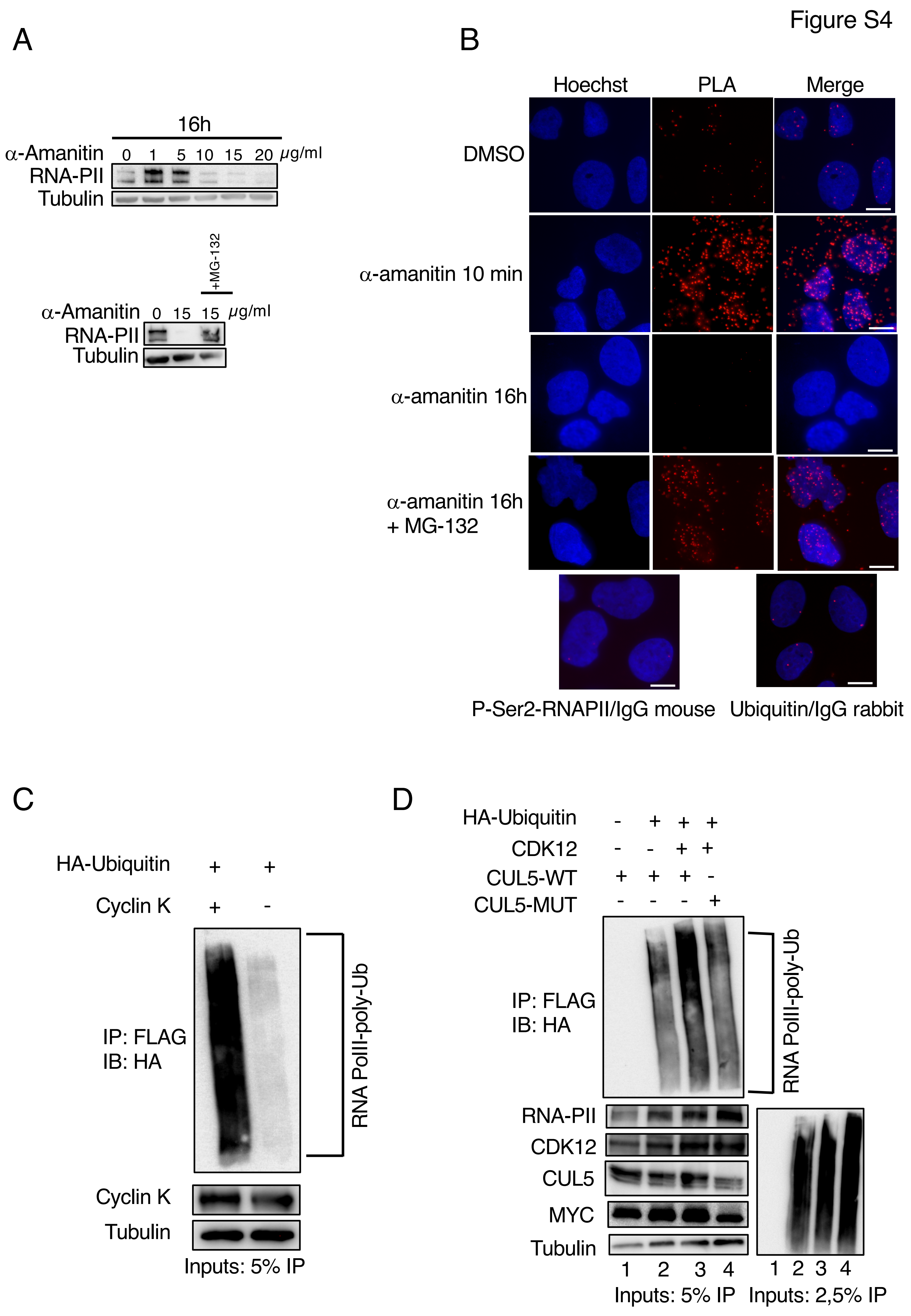

Figure S5

A

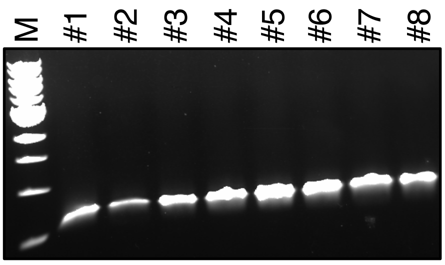


B

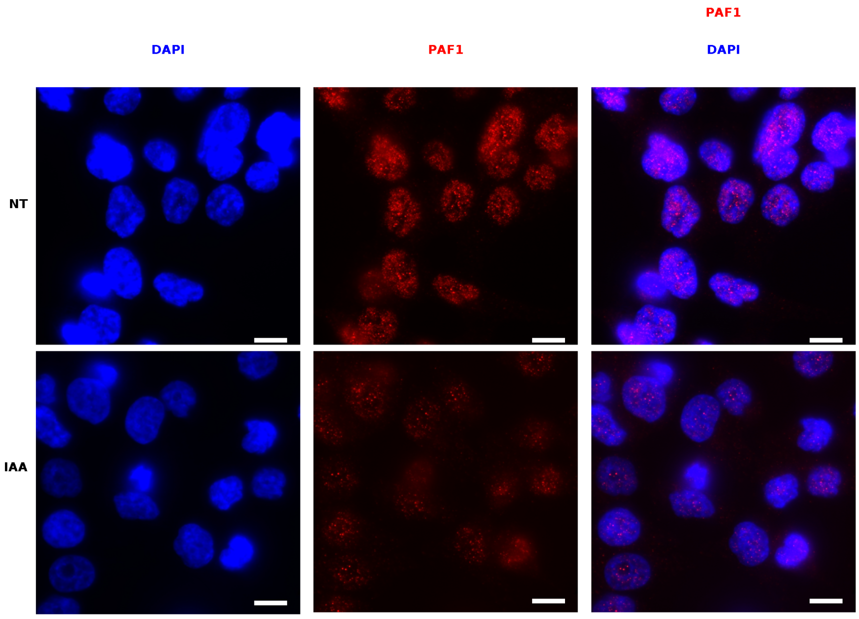


C

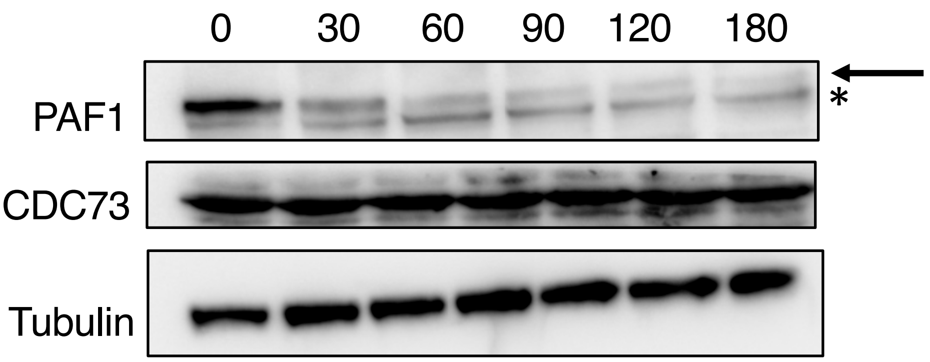


D

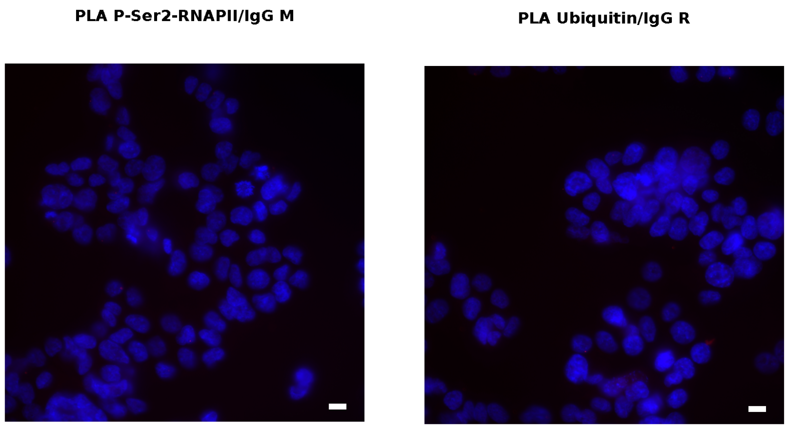
