## Supplemental Table S1 for "PAF1 facilitates RNA polymerase II ubiquitination by the Elongin A complex through phosphorylation by CDK12"

| reference | Gene Symbol | Unique | Total |
|---|---|---|---|
| LEO1_HUMAN | LEO1 | 54 | 396 |
| PAF1_HUMAN | PAF1 | 54 | 367 |
| CTR9_HUMAN | CTR9 | 68 | 345 |
| CDC73_HUMAN | CDC73 | 43 | 179 |
| WDR61_HUMAN | WDR61 | 11 | 31 |
| HSP7C_HUMAN | HSPA8 | 23 | 27 |
| DHX9_HUMAN | DHX9 | 23 | 23 |
| CAF1A_HUMAN | CHAF1A | 18 | 21 |
| PGAM5_HUMAN | PGAM5 | 11 | 19 |
| CAF1B_HUMAN | CHAF1B | 18 | 18 |
| HNRPU_HUMAN | HNRNPU | 15 | 16 |
| ELOA1_HUMAN | TCEB3 | 14 | 15 |
| TBA1A_HUMAN | TUBA1A | 13 | 15 |
| ROA2_HUMAN | HNRNPA2B1 | 12 | 15 |
| TR150_HUMAN | THRAP3 | 13 | 14 |
| GRP78_HUMAN | HSPA5 | 12 | 12 |
| DHX15_HUMAN | DHX15 | 12 | 12 |
| HS71L_HUMAN | HSPA1L | 9 | 12 |
| WDR82_HUMAN | WDR82 | 9 | 11 |
| BCLF1_HUMAN | BCLAF1 | 10 | 11 |
| TCPQ_HUMAN | CCT8 | 10 | 10 |
| SF3B3_HUMAN | SF3B3 | 10 | 10 |
| PP1RA_HUMAN | PPP1R10 | 10 | 10 |
| RFC1_HUMAN | RFC1 | 10 | 10 |
| NUCL_HUMAN | NCL | 10 | 10 |
| DDX21_HUMAN | DDX21 | 10 | 10 |
| TBB2A_HUMAN | TUBB2A | 9 | 9 |
| RS3_HUMAN | RPS3 | 9 | 9 |
| PR40A_HUMAN | PRPF40A | 9 | 9 |
| TCPB_HUMAN | CCT2 | 8 | 9 |
| ROA1_HUMAN | HNRNPA1 | 2 | 9 |
| RUVB2_HUMAN | RUVBL2 | 8 | 8 |
| NASP_HUMAN | NASP | 8 | 8 |
| TCPE_HUMAN | CCT5 | 8 | 8 |
| HSP71_HUMAN | HSPA1A | 7 | 8 |
| MCM2_HUMAN | MCM2 | 8 | 8 |
| RBBP4_HUMAN | RBBP4 | 6 | 8 |
| TCPA_HUMAN | TCP1 | 7 | 7 |
| RUVB1_HUMAN | RUVBL1 | 7 | 7 |
| HSP72_HUMAN | HSPA2 | 6 | 7 |
| TCPD_HUMAN | CCT4 | 6 | 6 |
| IPO4_HUMAN | IPO4 | 6 | 6 |
| NAT10_HUMAN | NAT10 | 6 | 6 |
| HNRPM_HUMAN | HNRNPM | 6 | 6 |
| GRP75_HUMAN | HSPA9 | 6 | 6 |
| SRRM1_HUMAN | SRRM1 | 5 | 6 |
| FUS_HUMAN | FUS | 5 | 6 |
| NOP56_HUMAN | NOP56 | 6 | 6 |
| ACTA_HUMAN | ACTA2 | 6 | 6 |
| HNRPD_HUMAN | HNRNPD | 4 | 6 |
| TCPG_HUMAN | CCT3 | 5 | 5 |
| TOP1_HUMAN | TOP1 | 5 | 5 |
| TCPH_HUMAN | CCT7 | 5 | 5 |
| FBRL_HUMAN | FBL | 5 | 5 |
| A2MG_HUMAN | A2M | 4 | 5 |
| TBB1_HUMAN | TUBB1 | 3 | 5 |
| RA1L2_HUMAN | HNRNPA1L2 | 5 | 5 |
| NPM_HUMAN | NPM1 | 5 | 5 |
| NOP58_HUMAN | NOP58 | 4 | 5 |
| TOX4_HUMAN | TOX4 | 4 | 4 |
| HNRH1_HUMAN | HNRNPH1 | 4 | 4 |
| TCPZ_HUMAN | CCT6A | 4 | 4 |
| RBM10_HUMAN | RBM10 | 4 | 4 |
| HNRPK_HUMAN | HNRNPK | 4 | 4 |
| ELOB_HUMAN | TCEB2 | 4 | 4 |

| | | | |
|---|---|---|---|
| ROA3_HUMAN | HNRNPA3 | 4 | 4 |
| SP16H_HUMAN | SUPT16H | 4 | 4 |
| ZBT24_HUMAN | ZBTB24 | 3 | 3 |
| TBB4A_HUMAN | TUBB4A | 3 | 3 |
| DNJC9_HUMAN | DNAJC9 | 3 | 3 |
| EF1A1_HUMAN | EEF1A1 | 3 | 3 |
| GPTC4_HUMAN | GPATCH4 | 3 | 3 |
| TBB3_HUMAN | TUBB3 | 3 | 3 |
| RRP1B_HUMAN | RRP1B | 3 | 3 |
| ASF1A_HUMAN | ASF1A | 3 | 3 |
| PCBP1_HUMAN | PCBP1 | 3 | 3 |
| MCM7_HUMAN | MCM7 | 3 | 3 |
| SMCA5_HUMAN | SMARCA5 | 3 | 3 |
| VIME_HUMAN | VIM | 3 | 3 |
| PRP4B_HUMAN | PRPF4B | 3 | 3 |
| H12_HUMAN | HIST1H1C | 3 | 3 |
| RL40_HUMAN | UBA52 | 3 | 3 |
| RS4X_HUMAN | RPS4X | 3 | 3 |
| TOX2_HUMAN | TOX2 | 2 | 2 |
| ROA0_HUMAN | HNRNPA0 | 2 | 2 |
| KEAP1_HUMAN | KEAP1 | 2 | 2 |
| SFPQ_HUMAN | SFPQ | 2 | 2 |
| RFC2_HUMAN | RFC2 | 2 | 2 |
| IgG1b_bovine | | 2 | 2 |
| RFC4_HUMAN | RFC4 | 2 | 2 |
| DDX5_HUMAN | DDX5 | 2 | 2 |
| NONO_HUMAN | NONO | 2 | 2 |
| RBM25_HUMAN | RBM25 | 2 | 2 |
| TBB5_HUMAN | TUBB | 2 | 2 |
| IMA2_HUMAN | KPNA2 | 2 | 2 |
| AKAP8_HUMAN | AKAP8 | 2 | 2 |
| H1X_HUMAN | H1FX | 2 | 2 |
| RL23_HUMAN | RPL23 | 2 | 2 |
| RL4_HUMAN | RPL4 | 2 | 2 |
| RS7_HUMAN | RPS7 | 2 | 2 |
| DDX18_HUMAN | DDX18 | 2 | 2 |
| MCM4_HUMAN | MCM4 | 2 | 2 |
| CSK21_HUMAN | CSNK2A1 | 2 | 2 |
| TOP1M_HUMAN | TOP1MT | 2 | 2 |
| HNRPF_HUMAN | HNRNPF | 2 | 2 |
| TRY3_HUMAN | PRSS3 | 1 | 2 |
| IgG1_bovine | | 1 | 2 |
| MCM6_HUMAN | MCM6 | 2 | 2 |
| H4_HUMAN | HIST1H4A | 2 | 2 |
| RL11_HUMAN | RPL11 | 2 | 2 |
| SSRP1_HUMAN | SSRP1 | 2 | 2 |
| ILF3_HUMAN | ILF3 | 2 | 2 |
| PIP_HUMAN | PIP | 2 | 2 |
| HSP76_HUMAN | HSPA6 | 1 | 2 |
| LC7L2_HUMAN | LUC7L2 | 1 | 1 |
| RNPS1_HUMAN | RNPS1 | 1 | 1 |
| SF3A1_HUMAN | SF3A1 | 1 | 1 |
| RS23_HUMAN | RPS23 | 1 | 1 |
| RS2_HUMAN | RPS2 | 1 | 1 |
| ILF2_HUMAN | ILF2 | 1 | 1 |
| MBB1A_HUMAN | MYBBP1A | 1 | 1 |
| RS15A_HUMAN | RPS15A | 1 | 1 |
| GLYR1_HUMAN | GLYR1 | 1 | 1 |
| PRP31_HUMAN | PRPF31 | 1 | 1 |
| SRRM2_HUMAN | SRRM2 | 1 | 1 |
| RFC5_HUMAN | RFC5 | 1 | 1 |
| MMTA2_HUMAN | MMTAG2 | 1 | 1 |
| IF4A1_HUMAN | EIF4A1 | 1 | 1 |
| UBR7_HUMAN | UBR7 | 1 | 1 |
| RS27L_HUMAN | RPS27L | 1 | 1 |
| RS10_HUMAN | RPS10 | 1 | 1 |

| | | | |
|---|---|---|---|
| CHERP_HUMAN | CHERP | 1 | 1 |
| ANM1_HUMAN | PRMT1 | 1 | 1 |
| PCBP2_HUMAN | PCBP2 | 1 | 1 |
| ANM5_HUMAN | PRMT5 | 1 | 1 |
| RS14_HUMAN | RPS14 | 1 | 1 |
| CP088_HUMAN | C16orf88 | 1 | 1 |
| ACINU_HUMAN | ACIN1 | 1 | 1 |
| DESM_HUMAN | DES | 1 | 1 |
| TBB4B_HUMAN | TUBB4B | 1 | 1 |
| TCPW_HUMAN | CCT6B | 1 | 1 |
| RL23A_HUMAN | RPL23A | 1 | 1 |
| MMS22_HUMAN | MMS22L | 1 | 1 |
| RL3_HUMAN | RPL3 | 1 | 1 |
| LMO7_HUMAN | LMO7 | 1 | 1 |
| U520_HUMAN | SNRNP200 | 1 | 1 |
| ELAV1_HUMAN | ELAVL1 | 1 | 1 |
| RS25_HUMAN | RPS25 | 1 | 1 |
| RL24_HUMAN | RPL24 | 1 | 1 |
| RS16_HUMAN | RPS16 | 1 | 1 |
| CHD3_HUMAN | CHD3 | 1 | 1 |
| RL15_HUMAN | RPL15 | 1 | 1 |
| RL12_HUMAN | RPL12 | 1 | 1 |
| RBM39_HUMAN | RBM39 | 1 | 1 |
| CO3_HUMAN | C3 | 1 | 1 |
| RS8_HUMAN | RPS8 | 1 | 1 |
| RL17_HUMAN | RPL17 | 1 | 1 |
| RL13_HUMAN | RPL13 | 1 | 1 |
| NUMA1_HUMAN | NUMA1 | 1 | 1 |
| MATR3_HUMAN | MATR3 | 1 | 1 |
| PYR1_HUMAN | CAD | 1 | 1 |
| NAKD1_HUMAN | NADKD1 | 1 | 1 |
| H2B1A_HUMAN | HIST1H2BA | 1 | 1 |
| REV1_HUMAN | REV1 | 1 | 1 |
| DYH2_HUMAN | DNAH2 | 1 | 1 |
| DDX54_HUMAN | DDX54 | 1 | 1 |
| KHDR1_HUMAN | KHDRBS1 | 1 | 1 |
| XP32_HUMAN | XP32 | 1 | 1 |
| ACTB_HUMAN | ACTB | 1 | 1 |
| RL8_HUMAN | RPL8 | 1 | 1 |
| CDSN_HUMAN | CDSN | 1 | 1 |
| GGCT_HUMAN | GGCT | 1 | 1 |
| H11_HUMAN | HIST1H1A | 1 | 1 |
| G3P_HUMAN | GAPDH | 1 | 1 |
| TOP2A_HUMAN | TOP2A | 1 | 1 |
| HNRDL_HUMAN | HNRPDL | 1 | 1 |
| KV201_HUMAN | | 1 | 1 |
| PRDX1_HUMAN | PRDX1 | 1 | 1 |
| IGLL5_HUMAN | IGLL5 | 1 | 1 |
| H31T_HUMAN | HIST3H3 | 1 | 1 |
| HBB_HUMAN | HBB | 1 | 1 |
| NOP14_HUMAN | NOP14 | 1 | 1 |
