## Supplemental Table S2 for "PAF1 facilitates RNA polymerase II ubiquitination by the Elongin A complex through phosphorylation by CDK12"

| Function | Genes |
|---|---|
| ATP catabolic process | DDX54\|RBBP4\|RUVBL2\|SMARCA5\|SNRNP200\|TOP2A |
| DNA conformation change | ACIN1\|ASF1A\|CDC73\|CHAF1A\|CHAF1B\|CTR9\|FBL\|HIST1H4A\|LEO1\|NPM1\|PAF1\|PRMT1\|RBBP4\|RUVBL1\|RUVBL2\|SF3B3\|SMARCA5\|TOP1\|TOP2A\|WDR61\|WDR82 |
| DNA-dependent DNA replication | MCM2\|MCM7\|MMS22L |
| DNA-templated transcription, initiation | BCLAF1\|SMARCA5\|THRAP3 |
| RNA processing | ACIN1\|CDC73\|CTR9\|DDX5\|DDX54\|HNRNPA2B1\|HNRNPF\|HNRNPH1\|HNRNPM\|HSPA1A\|LEO1\|PAF1\|PRMT5\|PRPF31\|RBM25\|RNPS1\|RPL11\|RPL14\|RPS16\|RPS7\|SF3A1\|SFPQ\|SNRNP200\|SRSF7\|THRAP3 |
| RNA stabilization | CDC73\|DHX9\|ELAVL1\|GAPDH\|HNRNPU\|KHDRBS1\|NPM1\|RPS14\|RPS27L\|RPS4X\|THRAP3\|YBX1 |
| cellular response to reactive oxygen species | CAT\|HBB\|PRDX1\|PRDX2\|TPM1 |
| erythrocyte differentiation | ACIN1\|HSPA1A\|RPS14 |
| intrinsic apoptotic signaling pathway by p53 class mediator | DDX5\|MYBBP1A\|RPS27L |
| myeloid leukocyte differentiation | ACIN1\|ANXA2\|CBFA2T3 |
| regulation of mRNA processing | ACIN1\|ANXA2\|ASF1A\|CBFA2T3\|CDC73\|CHAF1A\|CHAF1B\|CTR9\|DDX5\|FBL\|HIST1H4A\|HSPA1A\|LEO1\|NPM1\|PAF1\|PRMT1\|PRMT5\|RBBP4\|RBM25\|RNPS1\|RPS14\|RUVBL1\|RUVBL2\|SF3B3\|SMARCA5\|SRSF7\|THRAP3\|TOP2A\|WDR61\|WDR82 |
| retina homeostasis | ACTB\|AZGP1\|PIP\|PRDX1 |
| ribosome biogenesis | PRMT5\|PRPF31\|RPL11\|RPL14\|RPS14\|RPS16\|RPS7 |
