## Supplemental Table S3 for "PAF1 facilitates RNA polymerase II ubiquitination by the Elongin A complex through phosphorylation by CDK12"

| Function | Genes |
|---|---|
| Adherens junction | ACTB\|CSNK2A1\|LMO7 |
| Antiviral mechanism by IFN-stimulated genes | EIF4A1\|KPNA2\|UBA52 |
| Arrhythmogenic right ventricular cardiomyopathy (ARVC) | ACTA2\|ACTB\|DES\|LMNA\|TPM1\|VIM |
| Chromosome Maintenance | ACTB\|CSNK2A1\|HIST1H2AA\|HIST1H2BA\|HIST1H4A\|HIST3H3\|HSPA2\|LMNA\|MCM2\|MCM4\|MCM6\|MCM7\|NPM1\|NUMA1\|RBBP4\|REV1\|RFC1\|RFC2\|RFC4\|RFC5\|RUVBL1\|RUVBL2\|SMARCA5\|TUBA1A\|TUBB\|TUBB4A\|TUBB4B\|UBA52 |
| DNA strand elongation | CSNK2A1\|HIST1H2BA\|HIST1H4A\|HIST3H3\|HSPA2\|LMNA\|MCM2\|MCM4\|MCM6\|MCM7\|NPM1\|NUMA1\|RBBP4\|RFC1\|RFC2\|RFC4\|RFC5\|RUVBL1\|RUVBL2\|SMARCA5\|TUBA1A\|TUBB\|TUBB4A\|TUBB4B\|UBA52 |
| Destabilization of mRNA by AUF1 (hnRNP D0) | ELAVL1\|HNRNPD\|HSPA8\|UBA52 |
| Detoxification of Reactive Oxygen Species | CAT\|PRDX1\|PRDX2 |
| Formation of tubulin folding intermediates by CCT/TriC | ACTB\|C3\|CCT2\|CCT3\|CCT4\|CCT5\|CCT6A\|CCT7\|CCT8\|CSNK2A1\|EEF1A1\|EIF4A1\|HIST1H2BA\|HIST1H4A\|HIST3H3\|HSPA5\|HSPA9\|LMNA\|MCM2\|MCM4\|MCM6\|MCM7\|NCL\|NOP56\|NUMA1\|RBBP4\|RFC1\|RFC2\|RFC4\|RFC5\|RPL11\|RPL12\|RPL13\|RPL14\|RPL15\|RPL17\|RPL23\|RPL23A\|RPL24\|RPL3\|RPL4\|RPL8\|RPS10\|RPS14\|RPS15A\|RPS16\|RPS2\|RPS23\|RPS25\|RPS3\|RPS4X\|RPS7\|RPS8\|TCP1\|TUBA1A\|TUBB\|TUBB1\|TUBB2A\|TUBB3\|TUBB4A\|TUBB4B\|UBA52 |
| Legionellosis | ACTB\|C3\|CSNK2A1\|EEF1A1\|HSPA1A\|HSPA1L\|HSPA2\|HSPA5\|HSPA6\|HSPA8\|KPNA2\|PRSS3 |
| Pausing and recovery of Tat-mediated HIV elongation | RNPS1\|SRRM1\|SRSF3\|SRSF7\|SSRP1\|SUPT16H\|TCEB2\|TCEB3 |
| RNA Polymerase II Transcription | ACIN1\|RNPS1\|SRRM1\|SRSF3\|SRSF7\|SSRP1\|SUPT16H\|TCEB2\|TCEB3\|WDR82 |
| Ribosome | ACTB\|CCT2\|CCT3\|CCT4\|CCT5\|CCT6A\|CCT7\|CCT8\|CHD3\|DHX9\|EEF1A1\|EIF4A1\|ELAVL1\|FUS\|HIST1H2BA\|HIST1H4A\|HNRNPA0\|HNRNPA1\|HNRNPA2B1\|HNRNPA3\|HNRNPD\|HNRNPF\|HNRNPH1\|HNRNPK\|HNRNPM\|HNRNPU\|HSPA5\|HSPA8\|HSPA9\|LMNA\|NOP56\|PCBP1\|PCBP2\|PRMT5\|RBBP4\|RNPS1\|RPL11\|RPL12\|RPL13\|RPL14\|RPL15\|RPL17\|RPL23\|RPL23A\|RPL24\|RPL3\|RPL4\|RPL8\|RPS10\|RPS14\|RPS15A\|RPS16\|RPS2\|RPS23\|RPS25\|RPS27L\|RPS3\|RPS4X\|RPS7\|RPS8\|SF3A1\|SF3B3\|SNRNP200\|SRRM1\|SRSF3\|SRSF7\|SSRP1\|SUPT16H\|TCEB2\|TCEB3\|TCP1\|TUBA1A\|TUBB1\|TUBB2A\|TUBB3\|TUBB4A\|TUBB4B\|UBA52\|YBX1 |
