## Supplemental Table S4 for "PAF1 facilitates RNA polymerase II ubiquitination by the Elongin A complex through phosphorylation by CDK12"

| Antibody | Reference | Supplier | Application |
|---|---|---|---|
| Actin | A2066 | Sigma-Aldrich | WB |
| CCNK | NBP1-06519 | Novus Biologicals | IF, IP, WB |
| CDK12 | ABE1861 | Millipore | IP, WB |
| CDK12 | 26816-1-AP | ProteinTech | IP, WB |
| CUL5 | NBP1-22970 | Novus Biologicals | IP, WB |
| CUL5 | A302-173A | Bethyl laboratories | IP, WB |
| FLAG | M2 clone, F3166 | Sigma-Aldrich | IP, WB |
| HA | Beads-AC | SCBT | WB |
| LEO1 | A300-175A | Bethyl laboratories | IP, WB |
| MYC | 9E10, SAB4700447 | Sigma-Aldrich | WB |
| Mouse IgG<br>Rabbit IgG | 02-6502<br>02-6102 | ThermoFisher<br>ThermoFisher | IP<br>IP |
| PAF1 | NB600-273 | Novus Biologicals | WB |
| PAF1 | NB600-27 | Novus Biologicals | IP |
| PAF1 | Ab137519 | Abcam | IF |
| Phospho-Ser2-RNAPII | ab5095 | Abcam | IF, IP, WB,PLA |
| Phospho-Ser5-RNAPII | ab5131 | Abcam | WB |
| RNAPII | F12, sc-55492 | SCBT | WB |
| RNAPII | CTD4H8, 05-623 | Millipore | IF, IP, WB |
| TCEB2 (Elongin B) | A304-008A | Bethyl laboratories | WB |
| TCEB1(Elongin C) | A304-787A | Bethyl laboratories | WB |
| TCEB3 (Elongin A) | A-5, sc-373811 | SCBT | WB |
| TCEB3 (Elongin A) | | Bethyl laboratories | IP |
| Tubulin | DM1A clone, T6199 | Sigma-Aldrich | WB |
| Ubiquitin | FK2, AB120 | Tebu-Bio | IF, PLA |
| Ubiquitin, K48 linkage specific | A101-050 | Boston Biochem | WB |
| Ubiquitin, K63 linkage specific | HWA4C4 | Enzo Life Sciences | WB |
