## Supplemental Table S5 for "PAF1 facilitates RNA polymerase II ubiquitination by the Elongin A complex through phosphorylation by CDK12"

| Oligos | Sequence (5' to 3') |
|---|---|
| HA-N1Fb | TAAGCATCTAGACCCGTTTGCTAGTTAGTCCCG |
| HA-N1R | TGCTTAGGTACCAAGGTAGGAGGGGTGACAAATG |
| Inv-N1Fb | TAAGCATCTAGACCCGTTTGCTAGTTAGTCCCG |
| Inv-N1Fc | TAAGCAACTAGTGCGCCCACCATCCAGACCC |
| Inv-N1R | TGCTTAGTCGACAGCGACGAGGCGACAACAG |
| Nt-F1 | GTTGGGTCCTCAGCGTCTTG |
| Nt-R1 | TGTGGGGAGTGTCCAAATGC |
| mAID-F1 | TTCGTGAAGGTATCAATGGACGG |
| U6-F | GAGGGCCTATTTCCCATGATTCC |
| HA-seq1F | CCCGTTTGCTAGTTAGTCCCG |
| HA-seq1R | AAGGTAGGAGGGGTGACAAATG |
| pMK-Sal1-F | TAAGCAGTCGACGGTATCGATAAGCTTGATATCG |
| pMK-SpeI-R | TGCTTAACTAGTGCCTGCACCAGCGCCTTTATAC |
| pMK-EcoRI-R | TGCTTAGAATTCGCCTGCACCAGCGCCTTTATAC |
| MutCUL5 | AGCGGCAATTAGTAATGCTCAGCTG |
